# Modeling the Switching behavior of Functional Connectivity Microstates (FCμstates) as a Novel Biomarker for Mild Cognitive Impairment

**DOI:** 10.1101/619437

**Authors:** SI Dimitriadis, María Eugenia López, Fernando Maestu, Ernesto Pereda

## Abstract

It is evident the need for designing and validating novel biomarkers for the detection of mild cognitive impairment (MCI). MCI patients have a high risk of developing Alzheimer’s disease (AD), and for that reason the introduction of novel and reliable biomarkers is of significant clinical importance. Motivated by recent findings about the rich information of dynamic functional connectivity graphs (DFCGs) about brain (dys)function, we introduced a novel approach of identifying MCI based on magnetoencephalographic (MEG) resting state recordings.

The activity of different brain rhythms {δ, θ, α1, α2, β1, β2, γ1, γ2} was first beamformed with linear constrained minimum norm variance in the MEG data to determine ninety anatomical regions of interest (ROIs). A dynamic functional connectivity graph (DFCG) was then estimated using the imaginary part of phase lag value (iPLV) for both intra-frequency coupling (8) and also cross-frequency coupling pairs (28). We analyzed DFCG profiles of neuromagnetic resting state recordings of 18 Mild Cognitive Impairment (MCI) patients and 20 healthy controls. We followed our model of identifying the dominant intrinsic coupling mode (DICM) across MEG sources and temporal segments that further leads to the construction of an integrated DFCG (iDFCG). We then filtered statistically and topologically every snapshot of the iDFCG with data-driven approaches. Estimation of the normalized Laplacian transformation for every temporal segment of the iDFCG and the related eigenvalues created a 2D map based on the network metric time series of the eigenvalues (NMTSeigs). NMTSeigs preserves the non-stationarity of the fluctuated synchronizability of iDCFG for each subject. Employing the initial set of 20 healthy elders and 20 MCI patients, as training set, we built an overcomplete dictionary set of network microstates (nμstates). Afterward, we tested the whole procedure in an extra blind set of 20 subjects for external validation.

We succeeded a high classification accuracy on the blind dataset (85 %) which further supports the proposed Markovian modeling of the evolution of brain states. The adaptation of appropriate neuroinformatic tools that combine advanced signal processing and network neuroscience tools could manipulate properly the non-stationarity of time-resolved FC patterns revealing a robust biomarker for MCI.

## 1. Introduction

The major cause of clinical dementia in the elderly is that of Alzheimer’s type (DAT; Qiu et al., 2009), which is mainly characterized by loss of synapses, the accumulation of the Beta amyloid protein (Aβ) and the phosphorylation of the Tau protein. Due to the progressive loss of synapses, which alters the efficient communication within and between various brain subsystems, the DAT may be considered a disconnection syndrome (Delbeuck et al., 2003). The pathological changes of DAT start decades before the first clinical symptoms appear, thus it is important to design proper analytic pathways for analysing neuroimaging datasets via, e.g., the notion of brain connectivity, which allows an early detecting of such changes (Gomez et al., 2009; Stam et al., 2009; Maestu et al., 2015). It is extremely important to Alzheimer’s disease (AD) research to early identify preclinical and prodromal AD as it can assist clinical trials and targeted interventions (Livingston et al., 2017).

Mild cognitive impairment (MCI) is considered to be an intermediate clinical stage between the normal cognitive decline and DAT (Petersen and Negash, 2008). The main parts of the affected brain during the MCI, apart from those involved in action and thought, are those related to memory. For that reason, MCI patients face memory problems on a higher level compared to normal aged population but with no prevalent characteristic symptomatology of dementia-like reasoning or impaired judgement (Petersen et al., 2009). MCI is a heterogeneous state with different subtypes, which complicates in many cases the prediction to DAT (Portet et al., 2006). Additionally, it is also difficult to accurately discriminate symptomatic predementia (MCI) from healthy aging or dementia (DAT) (Petersen and Negash, 2008). Despite these difficulties to achieve an early diagnosis, an accurate identification of MCI should be attempted. Early diagnosis of MCI even in the absence of a healing strategy is significant for both pharmacological and non-pharmacological interventions. For that reason, new tools based on neuroimaging approaches are needed to increase the sensitivity in the detection of MCI.

Analysis of MEG recordings untangled the association between neural oscillations, functional connectivity assessment and neurophysiological activity (Brookes et al., 2011). Altered frequency-dependent functional patterns have been linked to the progression of cognitive decline (Poza et al., 2014). Alternative scenarios of analysing MEG recordings include to single channel analysis e.g. power analysis, functional connectivity and brain network analysis in resting state and also in task-based experiments (for a review see Mandal et al., 2018). Analysis of single channel recordings is a less complex approach that identified aberrant oscillations in AD primarily in the left temporal-parietal-occipital brain areas (Gomez et al., 2009). Functional connectivity (FC) and effective (EC) connectivity analysis revealed a loss of connectivity in AD compared to healthy control (HC) subjects found mostly in higher frequency bands (Gomez et al., 2017) while multiplex network analysis of MEG study in AD identified affected regions of hippocampus, posterior default mode network (DMN) and occipital areas (Yu et al., 2017). However, current clinical literature is limited and no strong conclusion can be drawn.

A recent multicenter MEG study addressed this issue using functional connectivity (FC) analysis (Maestu et al., 2015). It revealed a hypersynchronization in MCI as the most discriminative feature of brain connectivity, mainly over the fronto-parietal and inter-hemispheric links. This pattern was stable across the five different neuroimaging centres that participated in the study (Accuracy ~= 80 %), which might thereby be considered as a preclinical connectomic biomarker for MCI/DAT. Previous MEG studies based on connectivity analysis described a less organized functional brain network a hypersynchrony in fronto-parietal network in MCI subjects (Bajo et al., 2010; Buldú et al., 2011), while patients with DAT demonstrated a less synchronized brain network accompanied with a cognitive decline (Stam et al., 2009). This hypersynchronization might be a compensatory mechanism but it cannot be adaptive since patient’s network is closes compared to healthy olds to random (Buldú et al., 2011). In a recent MEG study comparing progressive MCI and stable MCI, authors described hypersynchronization in α band between anterior cingulate and posterior brain areas in the progressive MCI group (López et al., 2014.)

Spontaneous fluctuations of functional MRI (fMRI) blood-oxygen-level-dependent (BOLD) signals are temporally coherent between distinct spatial brain areas and not random. Biswal et al., (1995) demonstrated that fluctuations from motor areas were correlated even in the absence of a motor task. FC based on BOLD signal is modulated by cognitive and affective states (Richiardi et al., 2011 and Shirer et al., 2012), by learning (Bassett et al., 2011), and also spontaneously (Britz et al., 2010, Chang and Glover, 2010 and Kitzbichler et al., 2009).

When non-stationarity is taken into account and a dynamic functional connectivity (DFC) approach is adopted for studying FC patterns even in the absence of a task (resting-state), more sophisticated algorithmic analyses should be used. In this line, two studies have been recently published simultaneously that presented a data-driven methodology. In the first one, Allen et al. (2014) proposed a method based on *k*-means clustering, aimed to detect distinct ‘FC states’ in the resting brain. These authors clearly showed differences from the stationary static functional brain networks. The second study proposed a data-driven method focused on extracting, out of hundreds of functional connectivity graphs (FCGs) in a multi-trial experimental paradigm, distinct brain states called *functional connectivity microstates* (FCμstates; Dimitriadis et al., 2013a). Both approaches revealed the need of dynamic FC to explore brain dynamics via the notion of brain connectivity, as it is clear that brain FC ‘hops’ from one state to another (FCμstate) leading to a Markovian chain with characteristic favoured transitions between pairs of FCμstates (Allen et al., 2014; Dimitriadis et al., 2010a, 2013a,b).

In the last years, an increasing amount of human brain research based on functional imaging methods (electro-encephalography: EEG/ magnetoencephalography: MEG/ functional Magnetic Resonance Imaging: fMRI) have adopted a dynamic approach for exploring how brain connectivity fluctuates during resting-state and tasks alike (Allen et al., 2014; Bassett et al., 2011; Braun et al., 2015; Calhoun and Adali, 2016; Dimitriadis et al., 2010a, 2012a, 2013a,b, 2015a,b,c,d, 2016a,b; Ioannides et al., 2012; Laufs et al., 2003; Mantini et al., 2007; Chang and Glover, 2010; Handwerker et al., 2012; Liu and Duyn, 2013; Hutchison et al., 2013; Yang and Lin, 2015; Mylonas et al., 2015; Toppi et al., 2015; for reviews see Calhoun et al., 2014). The aforementioned studies have demonstrated the superiority of DFC as compared to a static connectivity analysis.

In parallel, the concept of cross-frequency coupling (CFC) is gaining attention lately in the neuroscience community, as evinced by the increasing number of papers published with the incorporation of this type of interaction in the analysis (Florin and Baillet, 2015; Tewarie et al., 2016; Antonakakis et al., 2016a,b; Dimitriadis et al., 2015c,d, 2016a,b; van Wijk and Fitzgerald, 2014). Specifically, intrinsic coupling modes and especially CFC bias the task-related response and are sensitive to various brain diseases and disorders such as DAT, Parkinson, etc. (see, e.g. Engel et al., 2013 for a review). More recent studies have shown that the dynamics of spontaneously generated neural activity can be informative about the functional organization of large-scale brain networks (Fox et al., 2005; He et al., 2008; Hipp et al., 2012), revealing intrinsically generated “coupling modes” at multiple spatial and temporal scales (Deco and Corbetta, 2011; Engel et al., 2013).

Based on the aforementioned methodological evidences in microscale, it is significant to explore the repertoire of intra and cross□frequency interactions across brain rhythms and brain areas under the same integrated graph model (Antonakakis et al., 2017; Dimitriadis et al., 2016b, 2017b, 2018a,c).

In a previous study, we demonstrated how to design a connectomic biomarker for MCI based on source-reconstructed MEG activity via static brain network analysis (Dimitriadis et al., 2018a). Here, we extended this work by proposing a scheme to design a dynamic connectomic biomarker under the framework of dynamic functional connectivity (DFC) analysis. Additionally, the proposed scheme will be validated in a second blind dataset (SID did not know anything about the labels).

For this aim, we analysed the MEG activity of healthy controls and MCI patients at resting-state (eyes-open) via DFC analysis. Based on a previous approach (Dimitriadis et al., 2016b; Dimitriadis et al., 2017b), we detected the dominant type of interaction per pair of MEG sources and temporal segment (Dimitriadis et al., 2018c). This approach produced a subject-specific dynamic functional connectivity graph (DFCG). This approach created a 2D matrix of size sources × temporal segments that describe the evolution of the eigenvalues across experimental time. Afterward, we used neural-gas to design overcomplete dictionaries for sparse representation of NMTS^eigen^ independently for the two groups (Dimitriadis et al., 2013a). Then, we validated the whole approach in a blind dataset to quantify the generalization of the proposed method.

In Materials and Methods section, we described the data acquisition, preprocessing steps, information about the datasets and the proposed methodological scheme. The Results section is devoted to describe the results including the prototypical network FCμstates, the accuracy of prediction in a blind dataset and network-based information of brain states. Finally, the Discussion section includes the discussion of the current research results with future extensions.

## 2. Material and Methods

### 2.1 Subjects and and Ethics Statement

The training dataset includes eighteen right-handed individuals with MCI (71.89 ± 4.51 years of age) and twenty-two age-and gender-matched neurologically intact controls (70.91 ± 3.85 years of age) were recorded. Table 1 summarizes their demographic characteristics. All participants were recruited from the Neurological Unit of the “The Hospital Universitario San Carlos”, Madrid, Spain. They were right handed (Olfield, 1971), and native Spanish speakers. We used also a set of twenty subjects of unknown label (blind author SD) for further validation of the proposed dynamic connectomic biomarker (DCB). Table 2 summarizes the mean and standard deviation of the demographic characteristics of controls and MCI subjects from the blind dataset. Including the blind subjects, the total sample consisted of 29 MCI and 31 controls. At the beginning, we used twenty subjects from each group to train the algorithm and we kept twenty (nine control subjects and eleven MCI) for blind classification.

**Table 1.** Mean ± standard deviation of the demographic characteristics of controls and MCIs. M= males; F= females; Educational level was grouped into five levels: 1) Illiterate, 2) Primary studies, 3) Elemental studies, 4) High school studies, and 5) University studies; MMSE= Mini-Mental State Examination; LH ICV, Left hippocampus normalized by total intracranial volume (ICV); RH ICV, Right hippocampus normalized by ICV.

|  | Age | Gender<br>(M/F) | Educational<br>level | MMSE | LH ICV | RH ICV |
| --- | --- | --- | --- | --- | --- | --- |
| **Control**<br>**(n=22)** | 70.91±3.853 | 9/13 | 3.50±1.225 | 29.32±0.646 | 0.0025±0.0003 | 0.0025±0.0003 |
| **MCI**<br>**(n=18)** | 71.89±4.510 | 7/11 | 2.71±1.359 | 27.24±1.954 | 0.0022±0.0005 | 0.0021±0.0005 |

**Table 2.** Mean ± standard deviation of the demographic characteristics of the blind sample of controls and MCIs. M= males; F= females; Educational level was grouped into five levels: 1) Illiterate, 2) Primary studies, 3) Elemental studies, 4) High school studies, and 5) University studies; MMSE= Mini-Mental State Examination; LH ICV, Left hippocampus normalized by total intracranial volume (ICV); RH ICV, Right hippocampus normalized by ICV

|  | Age | Gender<br>(M/F) | Educational<br>level | MMSE | LH ICV | RH ICV |
| --- | --- | --- | --- | --- | --- | --- |
| **Control**<br>**(n=9)** | 70.22±3.8333 | 1/8 | 3.44±1.333 | 29.33±0.707 | 0.0026±0.0005 | 0.0027±0.0004 |
| **MCI**<br>**(n=11)** | 73.45±3.297 | 7/4 | 3.91±1.221 | 26.90±2.132 | 0.0018±0.0004 | 0.0021±0.0004 |

To explore their cognitive and functional status, all participants were screened by means of a variety of standardized diagnostic instruments and underwent an extensive cognitive assessment, as described in López et al. (2016).

MCI diagnosis was established according to the National Institute on Aging-Alzheimer Association (NIA-AA) criteria (Albert et al., 2011), being all of them categorized as “MCI due to AD intermediate likelihood”. They met the clinical criteria and also presented hippocampal atrophy, which was measured by magnetic resonance (MRI). According to their cognitive profile, they were also classified as amnestic subtype (Petersen et al., 2001).

The whole sample was free of significant medical, neurological and/or psychiatric diseases (other than MCI). Exclusion criteria included: a modified Hachinski Ischemic score ≥4 (Rosen et al., 1980); a geriatric depression scale short-form score ≥5 (Yesavage et al., 1982); a T2-weighted MRI within 12 months before MEG screening with indication of infection, infarction, or focal lesions (rated by two independent experienced radiologists, Bai et al., 2012); and other possible causes of cognitive decline such as B_12_ deficit, diabetes mellitus, thyroid problems, syphilis, or human immunodeficiency virus (HIV). Finally, those participants with medical treatment that could affect the MEG activity (e.g., cholinesterase inhibitors) were required to interrupt it 48 h before the MEG recordings.

The present study was approved by the local ethics committee and all subjects signed an informed consent prior to their MEG recording.

### 2.2 MRI Acquisition and hippocampal volumes

3D T1 weighted anatomical brain magnetic MRI scans were collected with a General Electric 1.5 TMRI scanner, using a high resolution antenna and a homogenization PURE filter (Fast Spoiled Gradient Echo (FSPGR) sequence with parameters: TR/TE/TI = 11.2/4.2/ 450 ms; flip angle 12°; 1 mm slice thickness, a 256 × 256 matrix and FOV 25 cm). Freesurfer software (version 5.1.0.; (Fischl et al., 2002)) was used to obtain the hippocampal volumes, which were normalized with the overall intracranial volume (ICV) of each subject.

### 2.3 MEG Acquisition and preprocessing

Four minutes of eyes-open resting state data were recorded while the participants were seated in a 306-channel (one magnetometer and two orthogonal planar gradiometers per recording site, sampling frequency of 1kHz) Vectorview system (ElektaNeuromag) placed in a magnetically shielded room (VacuumSchmelze GmbH, Hanau, Germany) at the “Laboratory of Cognitive and Computational Neuroscience” (Madrid, Spain). Subjects had to fix their look in a cross, which was projected in a screen. The position of the head relative to the sensor array was monitored by four head position indicator (HPI) coils attached to the scalp (two on the mastoids and two on the forehead). These four coils along with the head shape of each subject (referenced to three anatomical fiducials: nasion, and left-right preauricular points) were acquired by using a three-dimensional Fastrak Polhemus system. Vertical ocular movements were monitored by two bipolar electrodes, which were placed above and below the left eye, and a third one to the earlobe, for electrical grounding.

Four head position indicator (HPI) coils were placed in the head of the subject, two in the forehead and two in the mastoids, in order to online estimate the head position. The HPI coils were fed during the whole acquisition, allowing for offline estimation of the head position.

Maxfilter software (version 2.2 Elekta Neuromag) was used to remove the external noise from the MEG data using the temporal extension of signal space separation (tSSS) with movement compensation (correlation threshold= 0,9m time window= 10s) (Taulu and Simola, 2006). This algorithm removes the signals whose origin is estimated outside the MEG helmet, while keeping intact the signals coming from inside the head. In addition, the continuous HPI acquisition, combined with the tSSS algorithm, allowed for the continuous movement compensation. As result, the signal used in the next steps comes from a set of virtual sensors whose position remains static respect to the head of the subject. Those subjects whose movement along the recording was larger than 25 mm were discarded, following the recommendations of the manufacturer.

### 2.4 Source Reconstruction

We generated a volumetric grid for the MNI template by adopting a homogenous separation of 1 cm in each direction, with one source placed in (0, 0, 0) in MNI coordinates. The whole procedure resulted in a source model with 2,459 sources inside the brain surface where each one consisted of three perpendicular dipoles. Every source was then labelled using the automated anatomical labelling (AAL) atlas (Tzourio-Mazoyer et al., 2002). We finally considered 1,467 cortical sources. The computed grid was then transformed to subject specific space employing the original T1 image. The realignment of grid and brain surface was realized manually to Neuromag coordinate system following the three fiducials and the head shape guides. Employing a realistically shaped head, we estimated a lead field (Nolte, 2003). We source reconstructed frequency-dependent brain activity using a Linearly Constrained Minimum Variance (LCMV) beamformer (Van Veen et al., 1997). We ran LCMV beamformer independently for the following eight frequency bands: δ (1-4 Hz), θ (4– 8 Hz), α_1_ (8–10 Hz), α_2_ (10-13 Hz), β_1_ (13–20 Hz), β_2_ (20-30 Hz), γ_1_ (30–49 Hz) and γ_2_ (51–90 Hz). The resulting spatial filters were projected over the maximal radial direction, getting only one spatial filter per source. “Radial direction” means the direction of the segment connecting the dipole location to the center of the sphere best approximating the brain surface. Radial dipoles in a spherical conductor do not produce a magnetic field outside of the conductor (Sarvas, 1987), so this projection avoids the creation of undetectable sources among the target dipoles. Finally, we represent every brain area – region of interest according to the AAL atlas by one source-space time series per frequency band using two alternative solutions: (1) the PCA of all the sources in the area or (2) the source closest to the centroid of the area (CENT).

Fig.1 illustrates the source-localization procedure and the different frequency-dependent representative Virtual Sensor time series for the two ROI representation schemes, PCA and the CENT.

**Figure 1.**
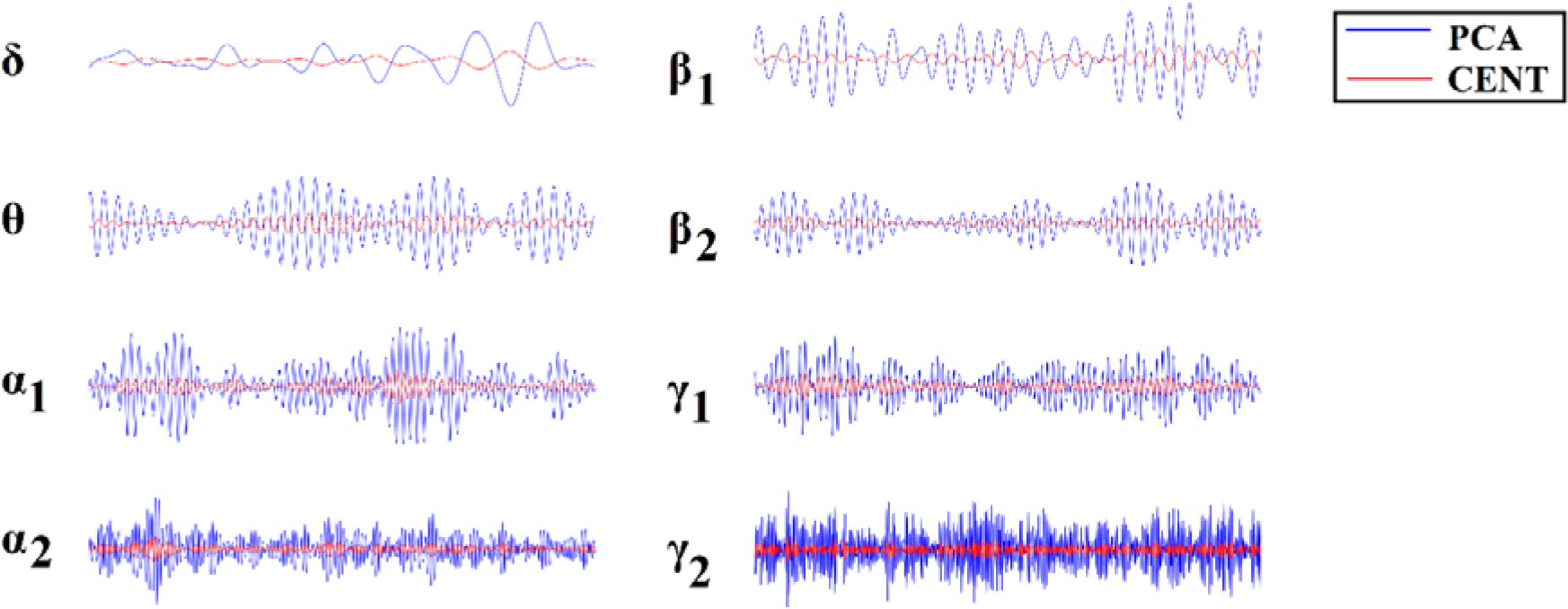
ROI Virtual Sensor representation of left precentral gyrus magnetoencephalographic activity from the first healthy control subject. Virtual sensor time series with blue and red color represent brain activity for PCA and CENT time series, respectively.

### 2.5 Dynamic Functional Connectivity Graphs (DFCGs)

#### 2.5.1 Construction of the integrated DFCGs

The DCFG analysis was restricted to the 90 ROIs of the AAL atlas. Adopting a common sliding window of width equal to 1s, to get at least 1 cycle of δ activity and a moving step of 50ms, we estimated the dynamic networks for both intra-frequency (8 frequency bands) and inter-frequency coupling modes (8*7/2=28 cross-frequency pairs) using the following formula of the imaginary part of phase locking value (iPLV).

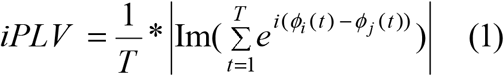

where *φ(t)*is the phase of the signal in the corresponding frequency band (intra-frequency modes) and between frequencies (cross-frequency couplings). For further details regarding phase-to-amplitude cross-frequency coupling (see Dimitriadis et al., 2015d and section1 in sup. material).

This procedure, whose implementation details can be found elsewhere (Dimitriadis et al., 2010a, 2015d, 2016b, 2017a, 2017b, 2018b), resulted in a four-dimensional tensor of size [coupling modes × temporal segments × ROIs × ROIs] or [36 × 2,401 × 90 × 90] time-varying PAC graphs per participant (^TV^PAC). Following proper surrogate analysis and a framework that have been presented in a previous study (Dimitriadis et al., 2018b), we defined the dominant intrinsic coupling mode (DICM) per pair of sources and across temporal segments. This procedure generates two three-dimensional tensors of size [temporal segments × ROIs × ROIs]. The first one keeps the functional coupling strength (iPLV) across anatomical space and time, while the second tabulates the DICM using an index for every possible case: {1 for δ, 2 for θ, 3 for α_1_, …,8 for γ_2_, 9 for δ-θ, …, 36 for γ_1_-γ_2_}. The following section described briefly the surrogate analysis appropriate for reducing pitfalls in CFC analysis and also to define the DICM.

#### 2.5.2 Statistical Filtering Scheme

At first place, we have to identify true CFC interactions that are not driven by the changes in signal power. Secondly, following a proper surrogate analysis our DICM model can detect the DICM between every pair of sources and at every temporal segment. The whole procedure of analysis is described in detail in (Dimitriadis et al., 2016b; Dimitriadis and Salis, 207b; Dimitriadis, 2018c). and also in see section 2 in sup.material

Fig.2 illustrates the whole procedure of DICM model for the first two temporal segments of resting-state activity of the first healthy control subject.

**Figure 2.**
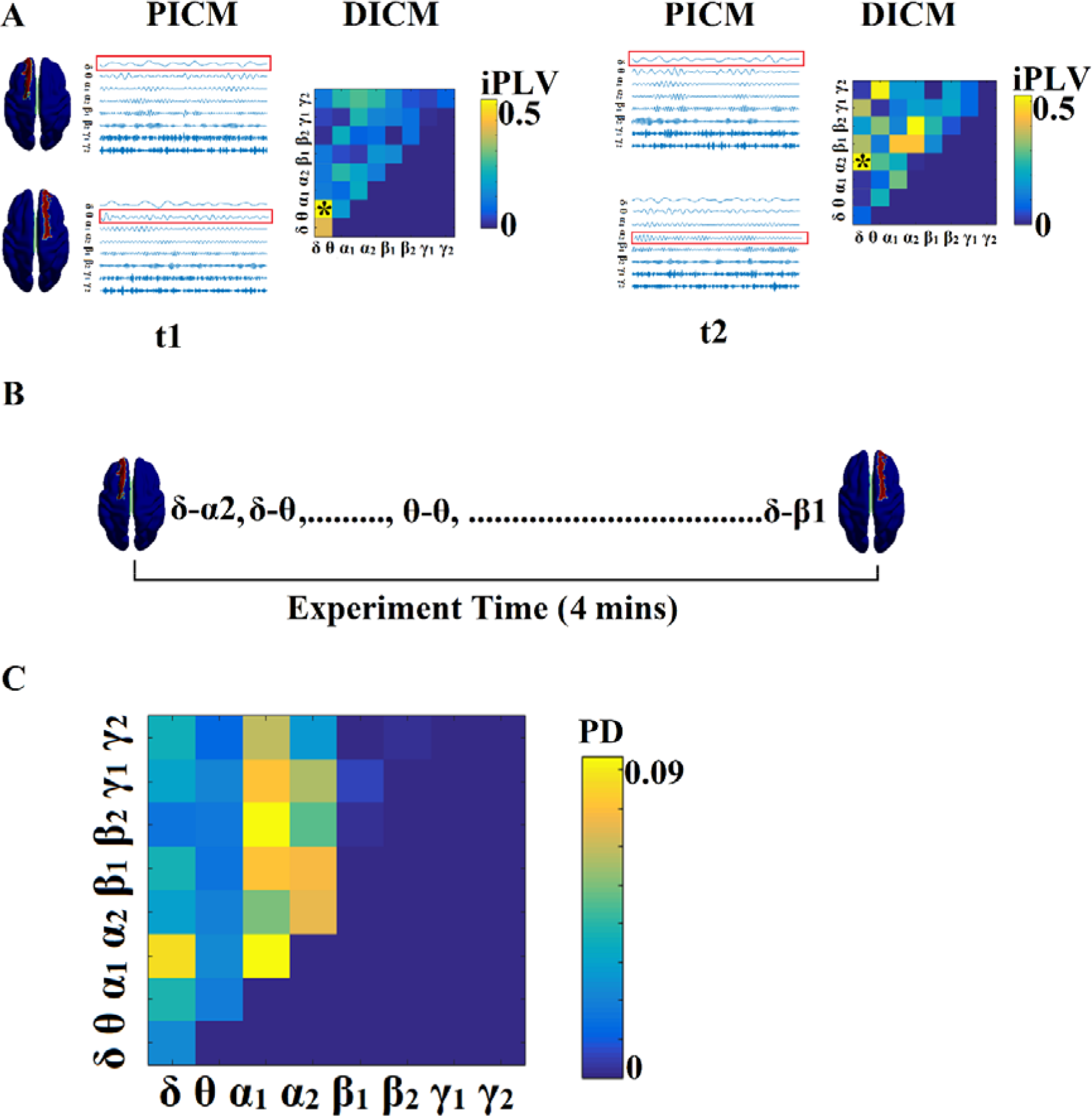
Determining Dominant Intrinsic Coupling Modes (DICM). An example for AU task derived from the first trial of first subject. A. Schematic illustration of our approach employed to identify the DICM between two sοurces (Left superior frontal gyrus, Right superior frontal gyrus) for two consecutive sliding time windows (t_1_, t_2_) during the first 4 secs of resting-state activity from the first healthy control subject. In this example, the functional synchronization between band-passed signals from the two sources was estimated by imaginary Phase Locking (iPLV). In this manner iPLV was computed between the two sources either for same-frequency oscillations (e.g., δ to δ…, γ_2_-γ_2_; 8 intra-frequency couplings) or between different frequencies (e.g., δ to θ, δ to α_1_…, γ_1_-γ_2_; 28 cross-frequency pairs). The sum of 8 + 28 = 36 refers to Potential Intrinsic Coupling Modes (PICM), which are tabulated in a matrix format. In the main diagonal, we inserted the intra-frequency couplings while in the off-diagonal the cross-frequency pairs. Statistical filtering, using surrogate data for reference, was employed to assess whether each iPLV value was significantly different from chance. During t_1_ the DICM reflected significant phase locking between α_1_ and α_2_ oscillations (indicated by red rectangles) in the oscillation list and a ‘*’ in the comodulogram. The DICM remains stable also for the t_2_ between α_1_ and α_2_ oscillations whereas during t_3_ the dominant interaction was detected between θ and α_2_ oscillations. B. Burst of DICM between Left and Right superior frontal gyrus. This packeting can be thought to associate the ‘letters’ contained in the DICM series to form a neural “word.”, a possible integration of many DICM. From this burst of DICM, we can quantify the probability distribution (PD) of DICM across experimental time (see C). C. Tabular representation of the probability distribution (PD) of DICM for left and right superior frontal gyrus across the experimental time shown in B. This matrix is called comodulogram and keeps the information of PD from the 36 possible coupling modes. In the main diagonal keeps the PD of the 8 possible intra-frequency coupling while in the off-diagonal the 28 possible cross-frequency pairs. (**Abbreviations: PICM**: Prominent Intrinsic Coupling Modes **DICM**: Dominant Intrinsic Coupling Modes **iPLV**:imaginary part of Phase Locking Value **PD**: probability distribution)

Fig.3.A demonstrates the first ten snapshots of the DFCG from the first healthy control subject.

**Figure 3.**
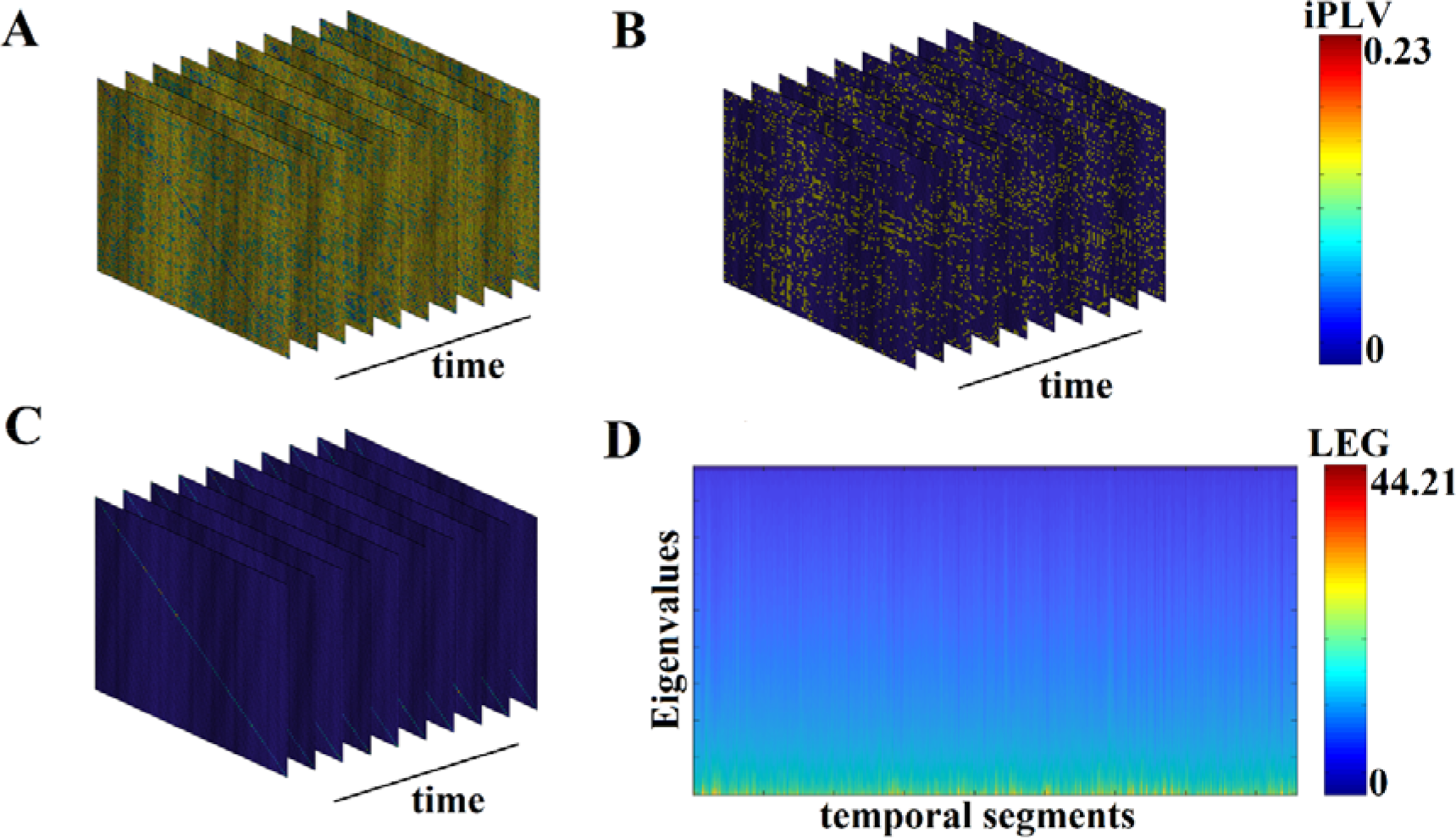
From DFCG to the temporal evolution of laplacian eigenvalues (LEG) (from the first healthy control subject). A) The very first ten snapshots of DFCG B) The quasi-static FCGs shown in A were topologically filtered with OMST C) The normalized laplacian transformation of the topologically filtered FCGs shown in B D) Temporal evolution of the laplacian eigenvalues for the 2.401 temporal segments

#### 2.5.3 Topological Filtering Scheme based on OMSTs

Apart from surrogate analysis, which is a statistical filtering procedure of the functional couplings within an FCG akin to a regularization to sparsify the 4D array described above, we adopted a topological filtering to further enhance the network topology and the most significant interactions. To this aim, we applied to each FCG derived from each subject and temporal segment independently, a novel data-driven thresholding scheme, proposed by our group, and termed Orthogonal Minimal Spanning Trees (OMSTs; Dimitriadis et al., 2017a, 2017c).

Fig.3.B demonstrates the temporal evolution of the topologically filtered dFCG for the first ten temporal segments.

#### 2.5.4 Graph Signal Processing

After extracting the most significant connections in DCFGs from each individual, we transformed every snapshot of the DFCG into the graph Laplacian variant called the normalized Laplacian matrix. With A being the functional connectivity graph and D, the degree matrix containing the degree of every node in the main diagonal, graph Laplacian L can be defined as L= D – A. The normalized graph Laplacian is defined as L_sym_ = D^−1/2^LD^−1/2^ (Shuman et al., 2013). We estimated the sorted eigenvalues of the Lsym for every snapshot of DFCG resulting to a 2-dimensional matrix of size [source (90) × temporal segments (2.401)] per subject. These 2D matrices were concatenated separately for the healthy control and disease group of the training set. Practically, the concatenation was performance along the temporal direction.

Fig.3.C shows the temporal evolution of the normalized Laplacian transformation of the dFCG for the first ten temporal segments while Fig.3.D is dedicated to the temporal evolution of the eigenvalues.

#### 2.5.5 A Vector-Quantization (VQ) Modeling of Group NMTS^eigen^

This subsection describes briefly our symbolization scheme, presented in greater details elsewhere (Dimitriadis et al., 2011, 2012, 2013a,b). The group-specific **NMTS^eigen^** patterns can be modelled as prototypical FC microstates (FCμstates). In our previous studies, we demonstrated a better modelling of DFCG based on vector quantization approach (Dimitriadis et al., 2013a, 2017b, 2018b). A codebook of *k* prototypical FC states (i.e., functional connectivity microstates-FCμstates) was first designed by applying the neural-gas algorithm (Dimitriadis et al., 2013a). This algorithm is an artificial neural network model, which converges efficiently to a small number *k* of codebook vectors, using a stochastic gradient descent procedure with a soft-max adaptation rule that minimizes the average distortion error (Martinetz et al., 1993). Neural-gas algorithm has been applied independently to each group by concatenating the 2D matrix of size [2.401 × 90] that describes the fluctuation of Laplacian eigenvalues.

The outcome of the neural-gas algorithm over NMTS^eigen^ is the construction of a symbolic sequence of group-specific prototypical FCμstates, one per subject. An example of such symbolic time series (STS) is a Markovian chain with three FCμstates: {1, 2, 3, 2, 1, 3, 2…} where each integer defines a unique brain state (FCμstates) assigned to every quasi-static temporal segment.

#### 2.5.6 External Validation in a blind dataset

We designed a novel approach of how to classify a blind subject. We reconstructed the subject-specific NMTS^eigen^ with both HC-based prototypical FCμstates and MCI-based prototypical FCμstates Specifically, for every temporal segment expressed via a vector of 90 eigenvalues, we estimated which of the prototypical FCμstates is much closer employing Euclidean distance for an appropriate criterion. Under this scheme, we rebuilt the original NMTS^eigen^ twice, one using prototypical FCμstates of HC and one using prototypical FCμstates of MCI. Then, we estimated the reconstruction mean squared error between the original NMTS^eigen^ and the two rebuilt NMTS^eigen^ based on prototypical FCμstates. Finally, we assign the test sample to the class with the lowest reconstruction error.

### 2.6 Markov chain modelling for synchro state transitions

The temporal sequence of spontaneous activity can be modelled as a Markovian process, which predicts the probabilities of several discrete states recurring or switching among themselves at different time points analysing time-point-based brain activity (Gärtner et al., 2015; Van de Ville, 2010). Several studies have investigated transition probabilities between phase-synchronized states on a sub-second temporal scale, untangling the Markovian property and the switching behaviour of finite network-level brain states (Baker et al., 2014; Dimitriadis et al., 2013,2015; Jamal et al., 2015).

#### 2.6.1 Markovian process of time-sequential FCμstates

Markov model describes the underlying dynamical nature of a system that follows a chain of linked states, where the appearance of a state at any given time instant depends only on the preceding ones (Gagniuc, 2017). In the Markov chain modelling for synchrostate transitions during the deductive reasoning and task-free processes, the first order transition matrices were estimated in a probabilistic framework. According to discrete-time Markov chain theory (Jarvis and Shier, 1999), a finite number (S_1_, S_2_…, S_m_) of inferred states, that evolve in discrete time with a time-homogeneous transition structure can be mathematically represented by either its -by-transition probability matrix or its directed graph (digraph). Here, the inferred states refer to the prototypical FCμstates. A feasible transition is one whose occurring probability is greater than zero. The probability of transition from node (state) *i* to node *j* is defined as

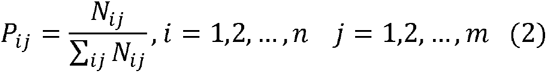

where *N*_*ij*_ is the number of transitions from node *i* to node *j*. Obviously, the sum of the transition probabilities along each row of the transition matrix P equals one. The complete digraph for a finite-state Markov process has edges of transition probabilities between every node i and every other node j. Here, nodes refer to FCμstates in the Markov chain. In the digraphs created in this study, if P_ij_ survives a p-value derived by 10,000 shuffled-surrogates of the original symbolic time series.

#### 2.6.2 Temporal measurements of an FCμstate symbolic sequence

For further summarizing inter-FCμstate transition patterns, relevant temporal measurements were obtained and analysed from the Markov chain structures of subject-specific FCμstate sequence, including: 1) fractional occupancy for each class of FCμstate (*i.e.*, the fraction of the number of distinct FCμstate of a given class occurring within 2.401 temporal segments), 2) dwell time for each FCμstate which gives the average time the brain spends within a specific FCμstate in consecutive temporal segments, 3) transition probabilities (TP) of a given FCμstate to any other functional connectivity state, 4) the complexity index (CI) that quantifies the richness of the spectrum of code-words formed up to a length based on the symbolic time series (Dimitriadis, 2018c) and 5) the flexibility index (FI) that quantifies the transition of the brainstates (FCμstates) between consecutive temporal segments.

#### 2.6.3 Assessing the Statistical Significant Level of the Symbolic-based Estimates

To assess the statistical significant level of the aforementioned four estimates (excluding CI), we shuffled the group symbolic time series 10,000 times and we re-estimated the surrogate-based p-values for every estimate per subject. CI is normalized by default with surrogates.

### 2.7 Linking MMSE with Chronnectomics

To investigate the possible relation between MMSE and the chronnectomics derived by the FCμstate symbolic sequence (see section 2.6.2), we used the canonical correlation analysis (CCA) approach to see whether MMSE correlates with seven chronnectomic variables.

### 2.7 Algorithms and MATLAB Code

All the algorithmic steps of constructing the DCFGs were implemented on in house software written in MATLAB, freely available from the first author’s github site https://github.com/stdimitr/from_dfcg_to_brain_states.

Same analysis is supported in PYTHON in a novel toolbox that van be found in the following link: https://github.com/makism/dyfunconn

LCMV beamformer has been programmed under Fieldtrip’s environment (Oostenveld et al., 2002).

## 3. Results

### 3.1 Group Prototypical FCμstates

Fig.4 illustrates the prototypical group-specific FCμstates for each group by assigning the 90 AAL brain areas to five well-known brain networks. The size and colour of every circle decodes the mean degree within every brain network while the colour of each connection defines the mean functional strength between every pair of brain networks. FCμstates can be described based on the most connected brain networks focusing on their degree. The most connected brain networks are the DMN and CO. Following a statistical test by comparing the functional coupling strength between FPN and DMN independently for every FCμstate, we found significant higher values for FCμstates 1 and 3 for HC compared to MCI (p = 0.00045 for FCμstate 1 and p=0.000012 for FCμstates 3, Wilcoxon Rank Sum Test).

**Fig.4.**
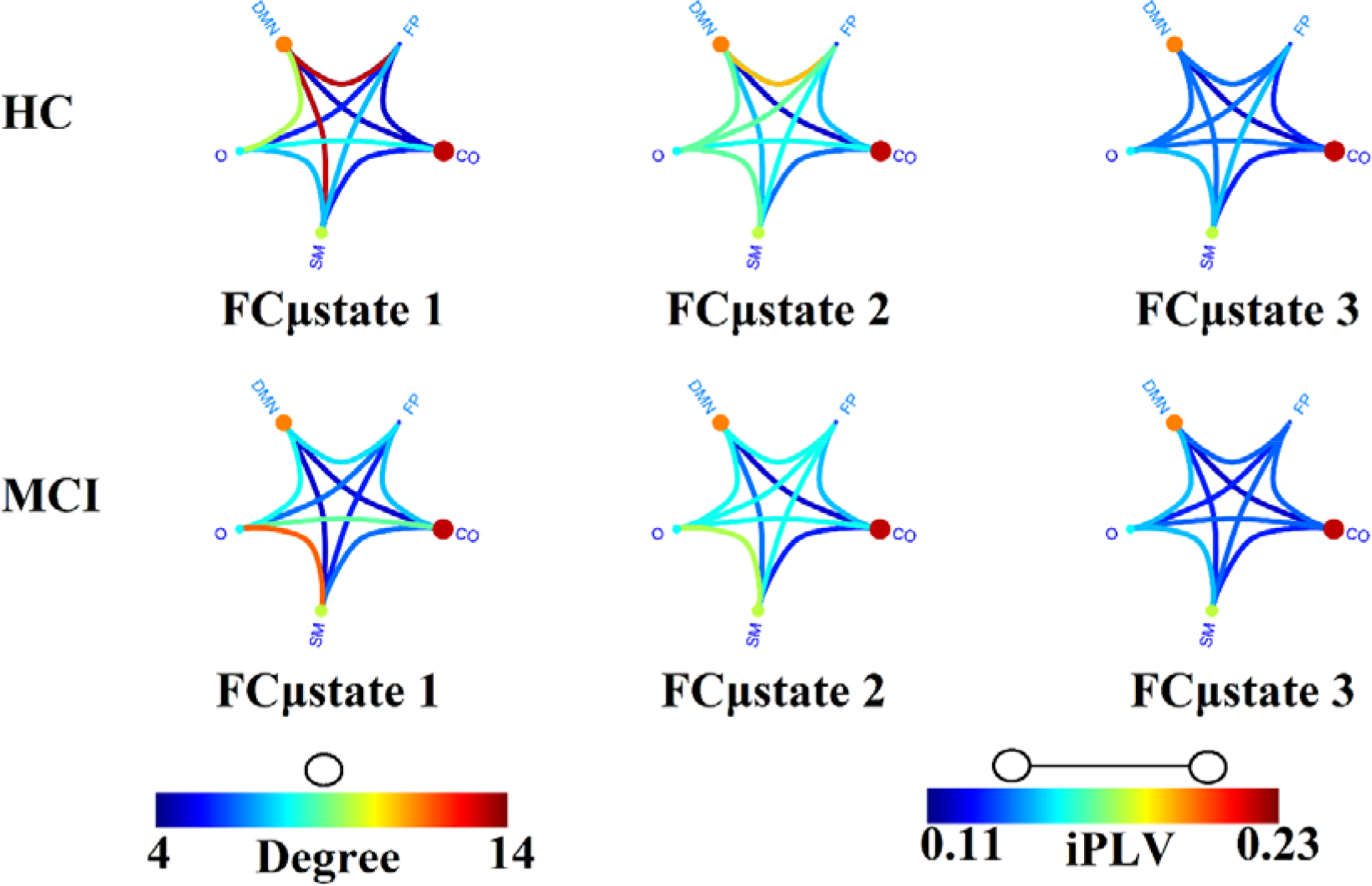
Prototypical FCμstates for healthy control (HC) and mild cognitive impairment (MCI). Neural-gas algorithm revealed three prototypical FCμstates per group with different spatial pattern. (FP: Fronto-Parietal, DMN: Default Mode Network, CO: Cingulo-Opercular, S: Sensorimotor, O: Occipital)

Fig.5 demonstrates the dynamic reconfiguration of prototypical FCμstates for the first subject of both groups for the first 1 min.

**Fig.5.**
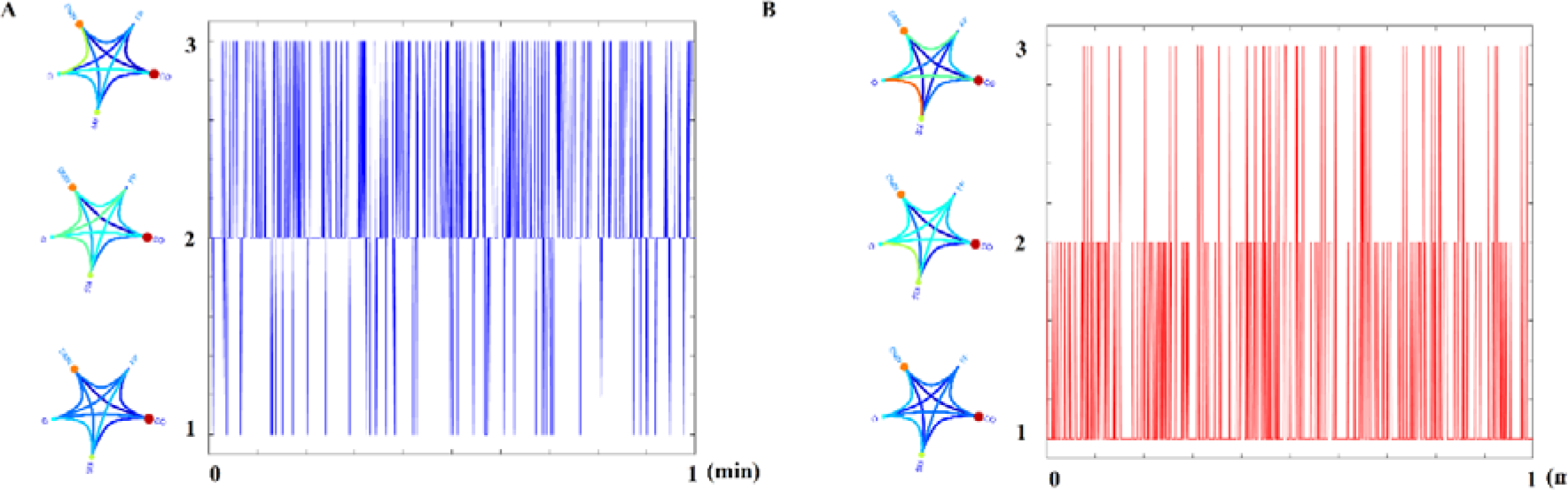
Temporal Evolution of Prototypical FCμstates for the first subject of A) HC and B) MCI group for the first min of resting-state.

### 3.2 Classification of Blind Samples via Representations with Prototypical Netμstates^eigen^

Each test sample with unknown label was classified to one of the two classes using as criterion the minimization of the reconstruction error. The minimum reconstruction error denotes the class label of the sample. In our study, we used twenty samples with distribution of 11 MCIs and 9 controls with 85% accuracy for CENT (17 out of 20) and 70% for PCA representation scheme (14 out of 20). SID received the blind dataset from MEL who evaluated the outcome of this research. Fig.6 illustrates the methodological approach. Fig.6A refers to the temporal resolution of the Laplacian eigenvalues of a blind HC subject while Fig.6B and Fig.6C the reconstruction of Fig.6.A matrix employing the **Prototypical Net**μ**states^eigen^** related to HC and MCI, correspondingly. Based on the reconstruction error between the original matrix (Fig.6A) and the two reconstructed (Fig.6B & Fig.6C), the decision about the label of the blind subject was taken based on the lowest reconstruction error.

**Figure 6.**
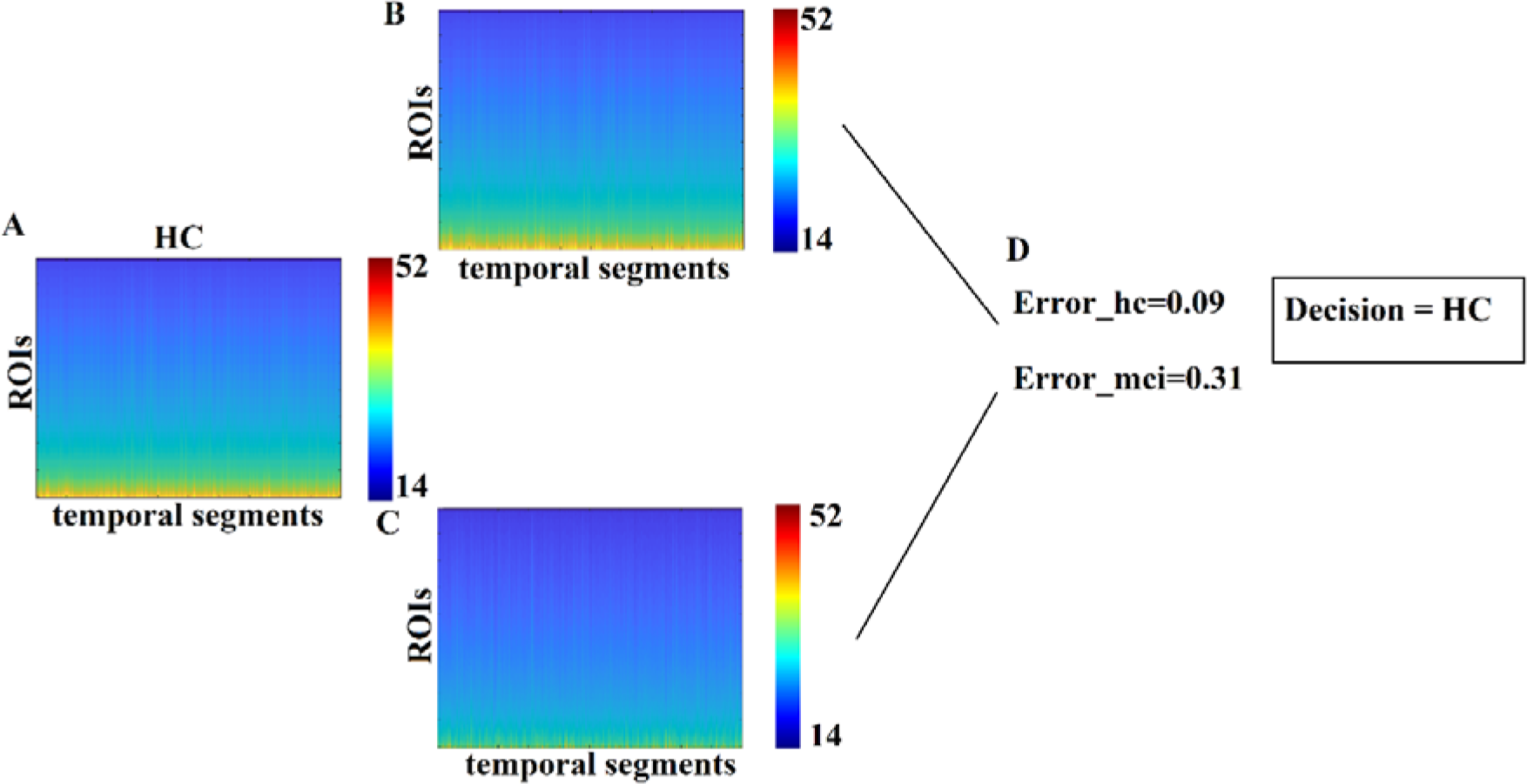
Classification of blind subjects via prototypical Netμstates^eigen^. **A.** Evolution of eigenvalues for the test sample **B.** Reconstruction of the temporal evolution of eigenvalues of the train sample with both group-specific prototypical Netμstates^eigen^ **C.** Estimation of the reconstruction error between the original temporal evolution of eigenvalues in A. and the two prototypical-based shown in B. The decision of subject’s label is taken via the lower reconstruction error.

### 3.3 Group-differences of Temporal measurements derived from FCμstate symbolic sequence

FI, OT and DT were significantly higher than the surrogates based FI values derived from the shuffled symbolic time series (p < 0.001). We detected significant higher FI and CI for HC compared to MCI applying a Wilcoxon Rank-Sum test (Fig.7, p-value < 0.00000001). Summarizing the results from OT and DT, HC subjects spent significantly higher time compared to MCI to first and third FCμstate while MCI spent significantly more time to the second FCμstate Fig.8, p-value < 0.00000001).

**Fig.7.**
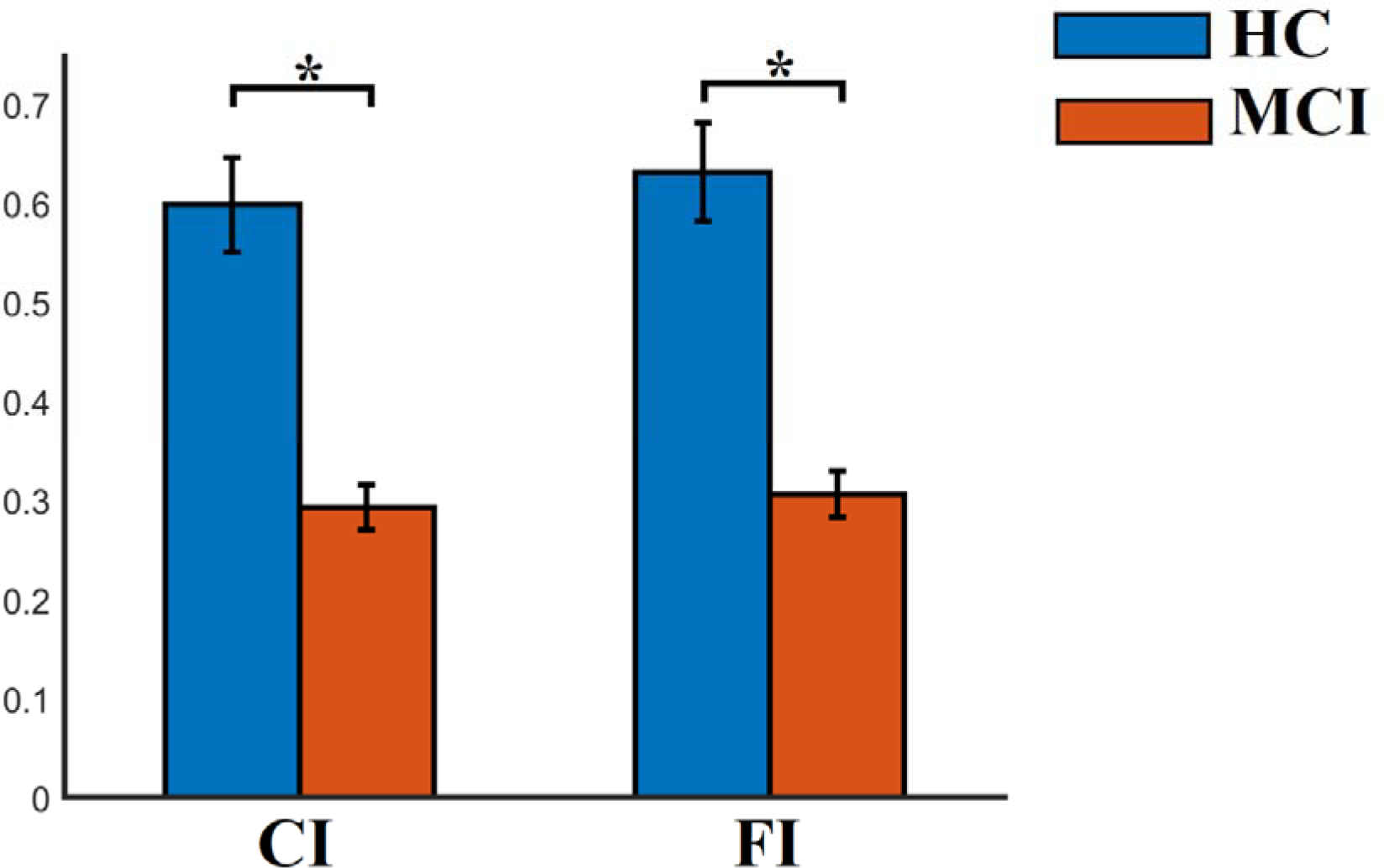
Group-averaged CI and FI. (* Wilcoxon Rank-Sum Test, p-value < 0.00000001)

**Fig.8.**
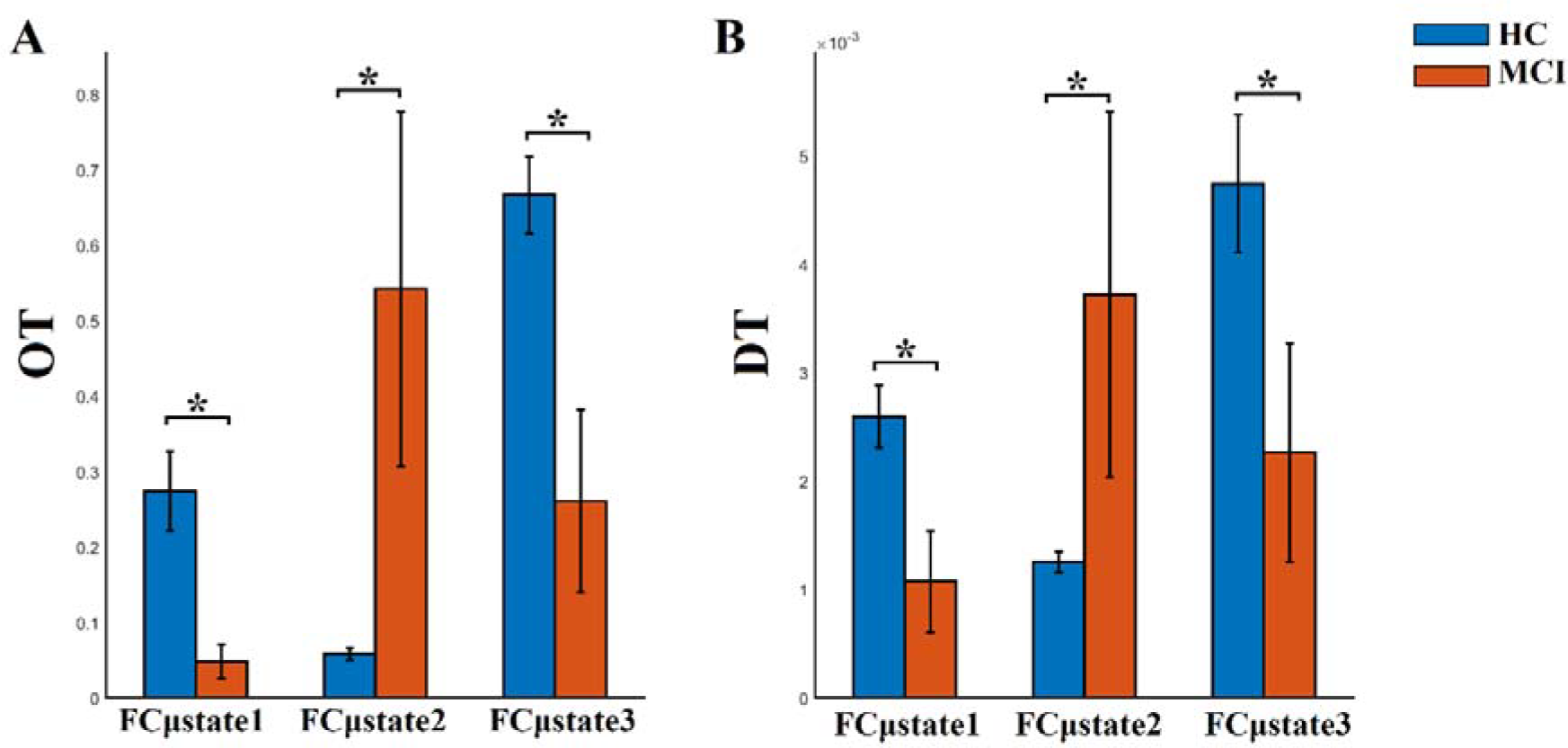
Group-averaged OT and DT per FCμstate. (* Wilcoxon Rank-Sum Test, p-value < 0.00000001)

### 3.4 Modelling Dynamic Reconfiguration of Functional Connectivity Graphs as a Markovian Chain

The outcome of the VQ modelling of NMTS^eigen^ is the derived Netμstates^eigen^ called FCμstates (see Fig.4) where its evolution is described via a symbolic time series, a Markovian chain. Fig. 9 illustrates a well-known scheme of the group-averaged transition probabilities (TP) between the three FCμstates for both groups. Our analysis revealed significant group-differences in terms of TP, while the TP were significant different compared to the surrogates’ symbolic time series. Self-loops defined the ‘staying’ TP of brain dynamics to the same brain state.

**Figure 9.**
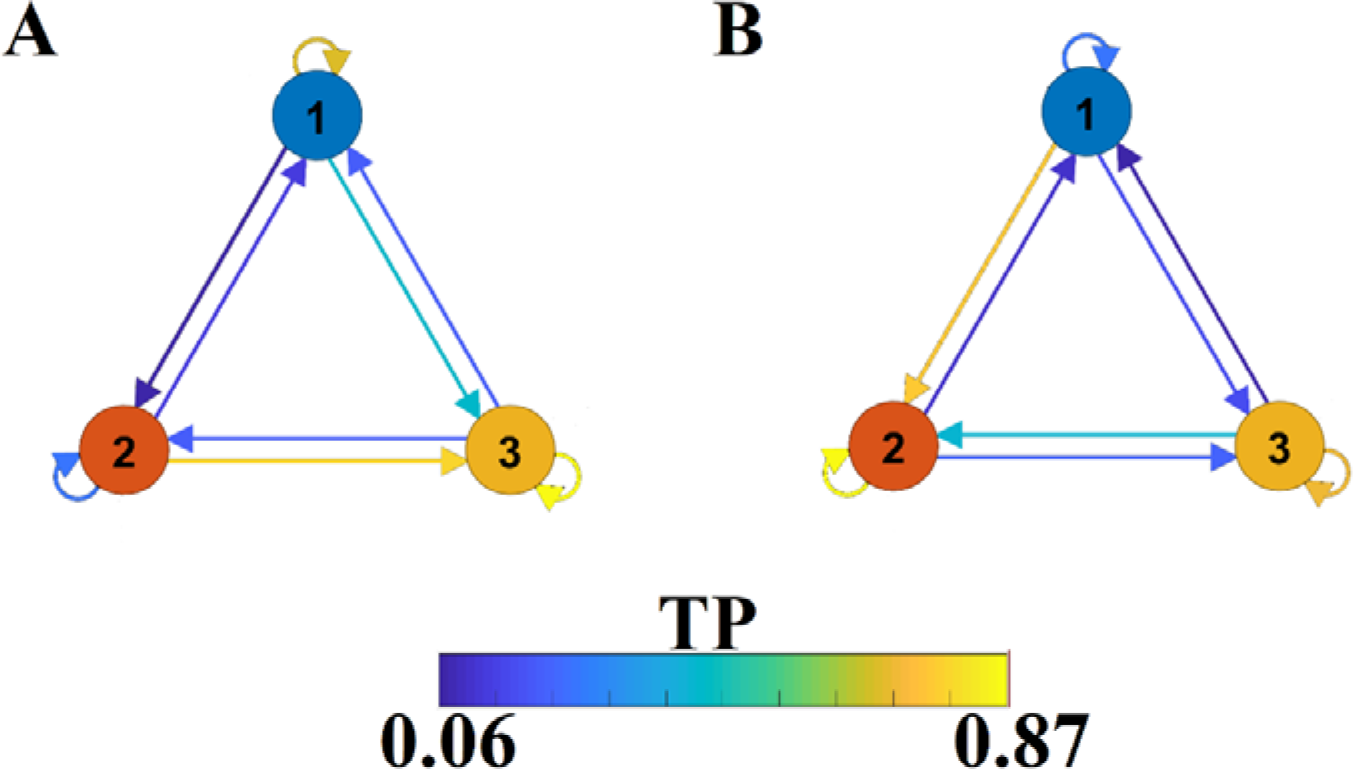
A finite-state diagram showing group-averaged transition probability matrix (TP) of the symbolic time series, which describes the temporal evolution of the brain, states (FCμstate). A) For healthy control and B) for mild cognitive impairment. The symbolic time series illustrated in Fig. 5 is a Markovian chain of order 1 and it is shown schematically with a diagram of three nodes defining the three FCμstates (Fig.4) while the arrows from one state to the other show the TP. Our results revealed significant group differences between every possible brain state transition (Wilcoxon Rank-Sum test, p < 0.0001/9).

### 3.5 Comodulograms of Dominant Intrinsic Coupling Modes (DICM)

Probability distributions (PD) of prominent intrinsic coupling modes across all sensors, sensor pairs, and time windows were summarized for each group in the form of an 8 × 8 matrix. The horizontal axis refers to the phase modulating frequency (Hz) while the y-axis to the amplitude modulated frequency (Hz). The main diagonal of the comodulograms keep the PD of intra-frequency phase-to-phase coupling. Group-averaged comodulograms in Figure 9 demonstrate the empirical PD of DICM revealing a significant role of α_1_ as phase modulator of the whole studying spectrum up to high-gamma (γ_2_) activity, which covers almost the 50% of pair-wise sensor connections and time-windows. No significant trend was detected regarding the PD of each pair of frequencies between the two groups (p < 0.05, Wilcoxon rank-sum test, Bonferroni corrected). Moreover, no significant difference was found regarding the PD of the groups for every possible pair of sensors (p < 0.05, Wilcoxon rank-sum test, Bonferroni corrected). Finally, transition dynamics of DICM between consecutive time-windows at every sensor-pair did not uncover any group-difference (for further details see Dimitriadis et al., 2016b).

**Figure 10.**
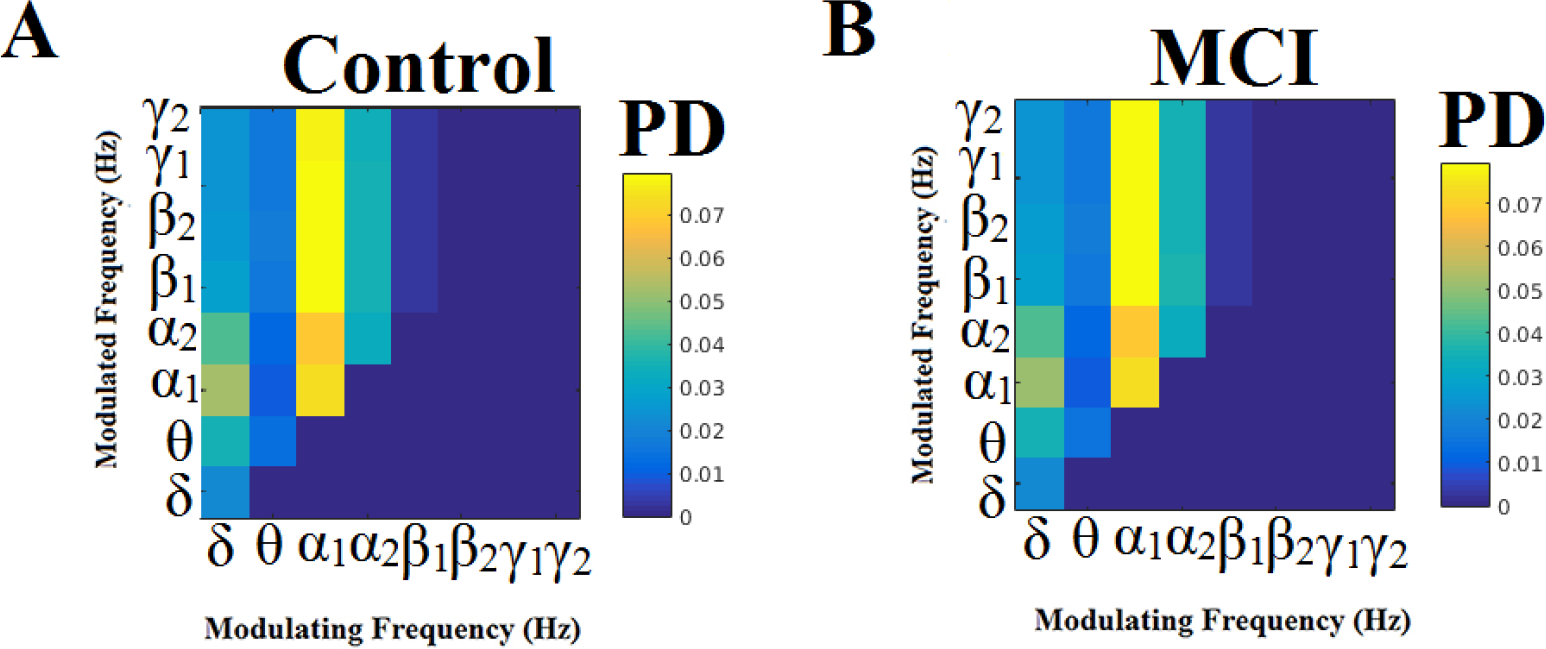
Group-averaged empirical Probability Distribution values of dominant intrinsic coupling modes for MCI (B) compared to control group (A).

### 3.6 Correlation of MMSE with Chronnectomics

Fig. 11 demonstrates the outcome of CCA analysis between chronnectomics and the well-known MMSE. The Chi-square was 26.95 and the related p-value = 0.00033886. X-axis refers to the canonical variable scores of the chronnectomics where DT of the three NMTS^eigen^ contributes most to the maximization of their canonical correlation with MMSE. OC_2 didn’t associate with the CCA mode of MMSE variability. The 1^st^ canonical component is: and the second is: CC2= 0.59*MMSE.

**Figure 11.**
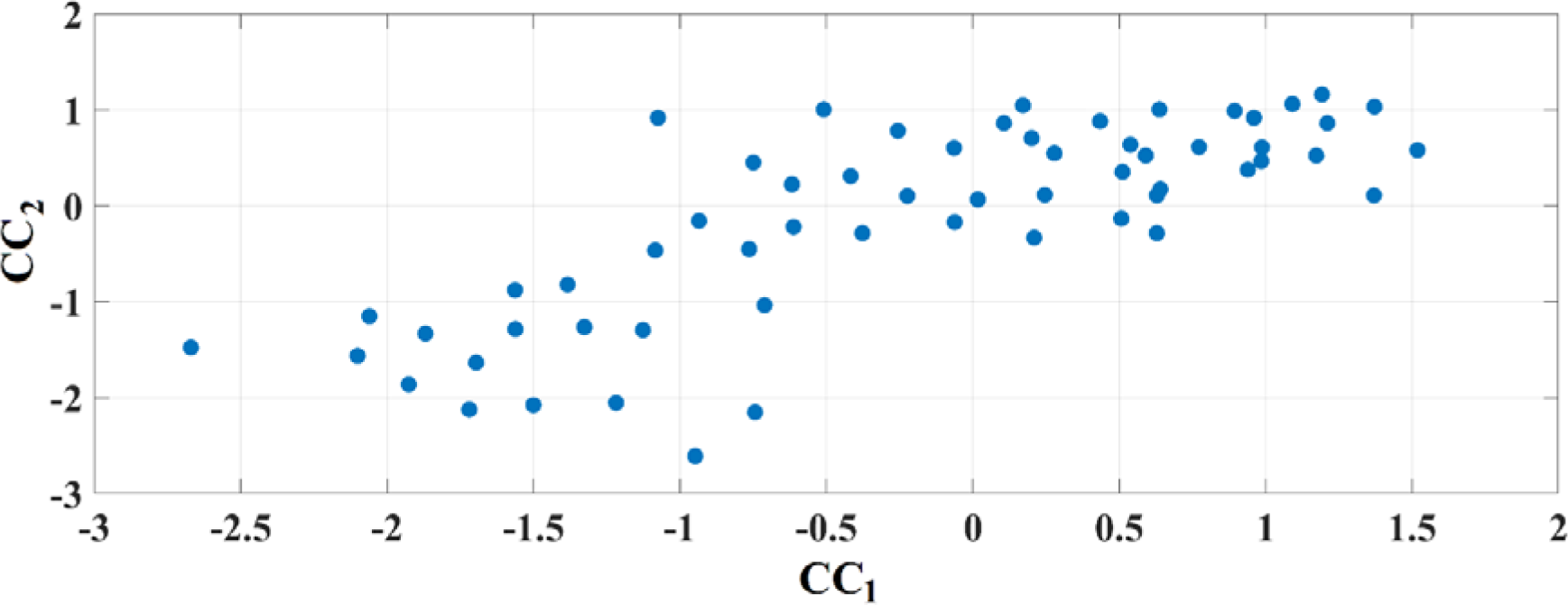
Canonical Correlation Analysis of Chronnectomics and MMSE. Figure plots the canonical variable scores referred to the two sets

## 4. Discussion

We have demonstrated here a novel framework for designing a proper dynamic connectomic biomarker (DCB) for the detection of MCI subjects from spontaneous neuromagnetic activity. The whole approach exhibits novel, data-driven, algorithmic steps that can be summarized as follows:

- The construction of a dynamic functional connectivity graph (DFCG) that incorporates dominant types of interactions, either intra-(e.g. θ-θ) or inter-frequency (phase-to-amplitude coupling (PAC) (e.g. θ-γ)) coupling.
- The application of a new thresholding scheme termed Orthogonal Minimal Spanning Trees (OMSTs) as a topological filtering applied to DFCG to extract a ‘true’ network topology.
- The VQ modelling of network metric time series (NMTS) based on nodal Laplacian eigenvalues for prototyping the spatiotemporal dynamics of both control and MCI subjects.
- Modelling of the switching behavior of brain states as a Markovian chain
- The validation of the whole approach to a second blind dataset achieving 85% of classification accuracy for the CENT ROI representation scheme compared to 70% for the PCA scheme
- ROI representation scheme matters on the designing of connectomic biomarker in general and also for MCI
- CCA analysis between chronnectomics and MMSE revealed that the DT of brain states associates strongly with the CCA mode of MMSE variability.

We proved that the VQ modelling of NMTS^eigen^ is an effective approach to extract an overcomplete dictionary for the representation of dynamic functional connectivity that can accurately classify subjects as either control or MCI based on their resting state MEG activity. Adopting a static network analysis, the classification accuracy was 12 out of 20, demonstrating the need of a dynamic functional connectivity approach for studying resting brain dynamics (Allen et al., 2014; Damarazu et al., 2014; Dimitriadis et al., 2010a, 2012c,d, 2013a,b, 2015b,d, 2016b, 2017a,b, 2018a,b; Kopell et al., 2014).

The capture of time-varying coupling between variables is a topic that has been heavily studied in other fields and in communications for signal processing in particular. However, the specific application to whole-brain functional connectivity is relatively new (Sakoglou et al., 2010; Dimitriadis et al., 2013a; Calhoun et al., 2014), and its application to brain-imaging data poses particular challenges, which are the topic of active current research. One important challenge is how to best identify relevant features from the high-dimensional brain imaging data. The main algorithms highly used for manipulating functional brain network dynamics in fMRI are group ICA (Calhoun and Adali, 2012) or spatial-constrained ICA (Lin et al., 2010) and tensor decompositions (Acar and Yener, 2009). Complementary, to characterize the dynamics of time-varying connectivity brain patterns, the basic approach is the metastate analysis based on the sliding window or more adaptive approach (Damaraju et al., 2015; Dimitriadis et al., 2013a; 2015a; Nomi et al., 2016). From the dynamic connectivity patterns, functional connectivity microstates (FCμstates) are extracted that are ‘quasi-stable’ distinct brain states. Then, the state vectors can be modelled via a Markovian chain (Calhoun et al., 2015; Damaraju et al., 2015; Dimitriadis et al., 2013a; 2015).

CFC mechanisms support the brain interactions across space over multiple temporal scales (Canolty & Knight, 2010; Fell & Axmacher, 2011). Computational models have explored the theoretical advantages of the existence of cross□frequency coupling (Lisman & Idiart, 1995; Neymotin et al., 2011). These models untangled the major mechanisms of the importance of CFC, which may serve as brain’s neural syntax. Segmentation of spike trains into cell clusters (‘letters’) and sequences of assemblies (neural ‘words’) is supported by the existing syntactic rules (Buzsaki, 2010).

In the present study, we demonstrated a methodology whose main scope is to provide a framework for modelling DFCG into a repertoire of distinct ‘quasi-static’ brain states called FCμstates. Here, we modelled the network metric time series (NMTS) derived by the dynamic functional connectivity patterns expressed via the Laplacian eigenvalues (Dimitriadis et al., 2015a, 2017b, 2018b). After extracting the virtual source time series, we followed an algorithmic approach with main aim to minimize the effect of a priori selection of variables that can minimize the reproducibility of the results. The main steps of the proposed methodology are: 1) the construction of one integrated dynamic functional connectivity per subject. which incorporates the dominant intrinsic coupling mode per each pair of brain areas and at every temporal segment, 2) the application of a data-driven topological filtering scheme to reveal the backbone of the network topology at every temporal segment, 3) the estimation of Laplacian eigenvalues to extract the so-called network metric time series (NMTS^eigen^; Dimitriadis et al., 2010, 2015a,2017b,2018b), 4) the modelling of these NMTS^eigen^ via a vector-quantization approach and 5) the validation of the whole approach to a second blind dataset.

The analysis of spatio-temporal evolution of Laplacian eigenvalues during the training phase revealed three prototypical brain states (FCμstates). For a better illustration of the FCGs linked to the prototypical eigenvalues, we assigned the 90 AAL brain areas to five well-known brain networks. In Fig. 4, we mapped the average functional strength between ROIs belonging to every pair of brain networks while the size and colour of every node defines the within brain network degree. The most connected brain networks in FCμstates are the DMN and CO. CO plays a key role in working memory mechanisms (Wallis et al., 2015) while cognitive complaints related to AD are linked to alterations of resting-state brain networks and mostly FPN and DMN (Contreras et al., 2017). Functional coupling strength between FPN and DMN was significantly higher for HC compared to MCI for FCμstates 1 and 3 (Fig. 4). The functional strength between FPN-DMN was positively correlated with a better episodic memory performance (Contreras et al., 2017).

Well-known and novel chronnectomics were estimated from the Markovian (symbolic) Chain that describes the evolution of brain states. We detected significant higher flexibility and complexity for HC as compared to MCI described from FI and CI, correspondingly (Fig.7). A summarization derived from OT and DT revealed a significant trend: HC subjects spent significantly higher time compared to MCI to FCμstates 1 and 3 while MCI spent significantly more time to the second FCμstate (Fig. 8). Following a CCA analysis between the extracted chronnectomics and the MMSE score; we found a significant contribution of the DT for the three NMTS^eigen^. OC related to the 2^nd^ NMTS^eigen^ didn’t associate with the CCA mode of MMSE variability (Fig.11).

In the era of data-sharing and aggregating large datasets from different research groups worldwide contributed to big consortiums, it is important to test the reproducibility of the proposed biomarkers (Abraham et al., 2016). Our study is a first step in this direction to diminish the effect of any arbitrary selection of algorithmic steps till the extraction of biomarkers. The next step is to reproduce the results on the source space and to extend the analysis in larger populations from different sites and MEG scanners. A recent study showed that a 70 percent of the preclinical research from academic labs could not be replicated (Collins and Tabak, 2014). Abraham’s work is one of the very first neuroimaging studies that lays the ground on the reliability and reproducibility of biomarkers extracted from neuroimaging data.

There is a large body of research based on different imaging methods covering various temporal and spatial scales that document the association of electrophysiological rhythms with distinct cognitive processes within narrowly or broadly anatomical areas (for review see Engel et al., 2001; Buszaki, 2006; Siegel et al., 2012; Bazar et al., 2013). For example, low frequency δ rhythms (1–4 Hz) are known to coordinate large portions of the brain (Fujisawa and Buzsaki, 2011; Nacher et al., 2013) while γ oscillations play a dominant role in stimulus processing and detection locally anatomically constrained (Engel et al., 2001). Recently, an extension of Brodmann’s areas was suggested in order to associate distinct anatomical areas with preferable connectivity estimator and cognitive functions in both normal and brain disease/disorder populations as an initial step to summarize the large body of current brain connectivity research (Basar et al., 2016).

In the last few years, an increasing number of studies appeared studying cross-frequency coupling (CFC) at resting-state (Antonakakis et al., 2016a), during cognitive tasks (Dimitriadis et al., 2015c,d, 2016a,b) and in various brain diseases and disorders such as mild traumatic brain injury (Antonakakis et al., 2016a), amnestic MCI (Dimitriadis et al., 2015d), dyslexia (Dimitriadis et al., 2016b), schizophrenia (Kirihara et al., 2012), etc. It has been suggested that CFC is the key mechanism for the integration of information between anatomically distribution subsystems that function on a dominant frequency (Canolty and Knight, 2010; Jirsa and Muller, 2013; Florin and Baillet, 2015). However, only a few MEG studies have explored CFC at resting-state (Antonakakis et al., 2015a, 2016 a,b; Florin and Baillet, 2015) and especially in a more dynamic fashion (Antonakakis et al., 2016b; Dimitriadis et al., 2015d, 2016b).

Our next goal is to work on the MEG source level after first improving sensitivity of beamformers in amplitude, phase and specifically to phase-to-amplitude coupling (PAC). MEG source connectivity is in a mature level compared to a decade ago (Ioannides et al., 2012), and it is an active research area aimed at improving many aspects of a ‘true’ brain connectivity (Schoffelen and Gross, 2009; Colclough et al., 2016). The most significant issue is the parcellation of cerebral cortex. In many cases, the AAL template (90 ROIs) is feasible for the detection of those changes induced by a specific task or obtained after compare different groups. But in others such as the design of a reliable connectomic biomarker, there is a need to oversample to more than 90 areas. The FC which is directly linked to functional parcellation of cerebral cortex is an active area, which will further improve both the interpretation and the predictive power of source connectivity of many brain diseases such as MCI. The solution of a functional parcellation template for MEG source connectivity will improve the classification performance on the source level with the additional advantage, compared to sensor level, of facilitating the anatomical interpretation of the results.

Adopting the same framework and including also stable and progressive MCI groups, we will attempt to connect DCB with neuropsychological measures and cognitive scores (Cuesta et al., 2015a,b). It is evident that a multifactorial model that includes cognition, neuropsychological measures and anatomical information can reliably predict the conversion from MCI to DAT, while genetic variation of risk genes like APOE-e4 allele or cognitive reserve might play a secondary role (Lopez et al., 2016).

Going one step further from our previous studies demonstrating the significance of a DCB (Dimitriadis et al., 2013b, 2015b), where we used network microstates extracted from DFCG patterns, in the present study we introduced a sparse modelling approach of NMTS^eigen^ estimated over DFCGs that preserve the dominant type of coupling (intra or inter-frequency intrinsic coupling mode). Our study demonstrates the effectiveness of the data-driven analytic pipeline tailored to DFCG to the correct classification of a blind dataset based on control and MCI subjects compared to a static connectivity approach. Given these outcomes, it is evident the need, in the next years, to adopt data-driven techniques that will not introduce bias, subjectiveness and assumptions in neuroimaging datasets and also to improve the reproducibility of the outcome in large databases.

In magnetoencephalography (MEG) the conventional approach to source reconstruction is to solve the underdetermined inverse problem independently over time and space. Different algorithms have been proposed so far with alternative regularization procedures of the space and time like with a Gaussian random field model (Solin et al., 2016).

Commonly used techniques include the minimum-norm estimate (MNE) (Hamalainen & Ilmoniemi, 1994) and Linearly Constrained Minimum-Variance (LCMV) beamformer (Van Veen et al., 1997). It is on the right direction to compare the consistency of the outcome of current study with alternative inverse solution algorithms to measure their consistency and sensitivity to the design of connectomic biomarkers tailored to MCI.

## 5. Conclusions

In this study, we presented a novel DCM for the prediction of MCI from age-matched control group validated over a blind dataset. The novelties of the proposed analytic scheme is the incorporation in the DFCGs the dominant intrinsic coupling mode (DICM, either intra or inter-frequency coupling based on PAC), the adaptation of a novel data-driven thresholding scheme based on OMSTs, the estimation of laplacian eigenvalues across time and the extraction of prototypical network microstates (FCμstates) for both control and MCI group.

It is important for the near future to work in source space on MCI subjects that convert to AD after a following up study to further validate the proposed scheme as a potential tool of clinical importance. Complementary, it would be interesting to explore how Apoe-e4 allele can induce changes to the DFC of spontaneous activity. Moreover, multimodal neuroimaging biomarkers is a novel trend that will further been validated (Jack et al., 2016).

## Acknowledgments

This study was supported by three projects from the Spanish Ministry of Economy and Competitiveness (PSI2009-14415-C03-01, PSI2012-38375-C03-01, TEC2016-80063-C3-2-R). This study was supported by the National Centre for Mental Health (NCMH) at Cardiff University. SID was supported by MRC grant MR/K004360/1 (Behavioural and Neurophysiological Effects of Schizophrenia Risk Genes: A Multi-locus, Pathway Based Approach). SD is also supported by a MARIE-CURIE COFUND EU-UK Research Fellowship. We would like to acknowledge RCUK of Cardiff University and Wellcome Trust for covering the publication fee.

## Author Contributions

SID: conception of the research analysis, methods and design, data analysis, and drafting the manuscript; MEL: data acquisition; MEL, FM, EP: critical revision of the manuscript. Every author read and approved the final version of the manuscript.

## Conflict of Interest Statement

The authors declare that the research was conducted in the absence of any commercial or financial relationships that could be construed as a potential conflict of interest.

## References

Abraham A, Milham M,Craddock RC, Samaras D, Thirion B, G Varoquaux.Deriving reproducible biomarkers from multi-site resting-state data: An Autism-based example. Neuroimage http://dx.doi.org/10.1016/j.neuroimage.2016.10.045

Acar E and B. Yener, 2009. Unsupervised multiway data analysis: A literature survey, *IEEE Trans*. Knowledge Data Eng., 21(1), pp. 6–20.

Aharon A, Elad M, Bruckstein A., 2006. K-SVD: An algorithm for designing overcomplete dictionaries for sparse representation. IEEE Transactions on Signal Processing 54(11), 4311–4322.

Albert, M. S., DeKosky, S. T., Dickson, D., Dubois, B., Feldman, H. H., Fox, N. C., et al., 2011. The diagnosis of mild cognitive impairment due to Alzheimer’s disease: recommendations from the National Institute on Aging-Alzheimer’s Association workgroups on diagnostic guidelines for Alzheimer’s disease. Alzheimers. Dement. 7, 270–9.

Allen E.A., Damaraju E., Plis S.M., Erhardt EB, Eichele T, Calhoun V.D., 2014. Tracking whole-brain connectivity dynamics in the resting state. Cereb Cortex. 24(3), 663–676.

Antonakakis, M., Dimitriadis SI, Zervakis M, Rezaie R, Babajani-Feremi A, Micheloyannis S, Zouridakis G, Papanicolaou A.C., 2015. Uncovering the brain model of MEG Brain Networks from Cross-Frequency Coupling Estimates via an attacking strategy. In Proceedings of the IEEE 37^th^ Int. Conf. on Engineering in Medicine and Biology Society.

Antonakakis, M., Dimitriadis, S.I., Zervakis M, Micheloyannis S, Rezaie R, Babajani-Feremi A, Gouridakis G, Papanicolaou A.C., 2016a. Altered cross-frequency coupling in resting-state MEG after mild traumatic brain injury. Int J Psychophysiol 102, 1–11.

Antonakakis,M., Dimitriadis S.I., Zervakis M, Zouridakis G, Papanicolaou A.C., 2016b. Mining Cross-Frequency Coupling Microstates from Resting State MEG: An Application to Mild Traumatic Brain Injury. In Proceedings of the IEEE 38^th^ Int. Conf. on Engineering in Medicine and Biology Society

Antonakakis M, Dimitriadis SI, Zervakis M, Papanicolaou AC, Zouridakis G.Altered Rich-Club and Frequency-Dependent Subnetwork Organization in Mild Traumatic Brain Injury: A MEG Resting-State Study. Front Hum Neurosci. 2017; 11:416. doi: 10.3389/fnhum.2017.00416

Aru, J., Priesemann, V., Wibral M., Lana L., Pipa G, Singer W, Vicente R., 2015. Untangling cross-frequency coupling in neuroscience. Curr Opin Neurobiol, 31, 51–61.

Bajo, R., Maestú, F., Nevado, A., Sancho M, Gutiérrez R, Campo P, Castellanos NP, Gil P, Moratti S, Pereda, E., 2010. Functional connectivity in mild cognitive impairment during a memory task: implications for the disconnection hypothesis. J. Alzheimers Dis. 22(1), 183–193.

Baker, A.P., Brookes, M.J., Rezek, I.A., Smith, S.M., Behrens, T., Probert Smith, P.J., et al., 2014.Fast transient networks in spontaneous human brain activity. Elife, 3(3), e01867.

Bai, F., Shu, N., Yuan, Y., Shi Y, Yu H, Wu D, Wang J, Xia M, He Y, Zhang Z (2012): Topologically convergent and divergent structural connectivity patterns between patients with remitted geriatric depression and amnestic mild cognitive impairment. J Neurosci 32, 4307–4318.

Bassett, D., Wymbs, N., Porter M, Mucha P, Carlson J, Grafton S., 2011. Dynamic reconfiguration of human brain networks during learning. Proc. Natl. Acad. Sci. U. S. A., 108, 7641–7646

Başar, E.B.G., 2013. Review of delta, theta, alpha, beta and gamma response oscillations in neuropsychiatric disorders. Suppl. Clin. Neurophysiol. 62,303–341

Başar E., Düzgün, A., 2016. The CLAIR model: Extention of Brodmann’s areas based on brain oscillations and connectivity. Int. J. Psychophysiol. 103,185–98.

Biswal, B., Yetkin, F.Z., Haughton V, M., Hyde, J.S., 1995. Functional connectivity in the motor cortex of resting human brain using echo-planar MRI Magn. Reson. Med., 34,537–54

Braun,U., Sarah F. Muldoon, Bassett D.S., 2014. On Human Brain Networks in Health and Disease.eLS (Wiley), DOI: 10.1002/9780470015902.a0025783.

Britz J.D., Van De Ville, C.M. Michel C.M., 2010. BOLD correlates of EEG topography reveal rapid resting-state network dynamics. Neuroimage, 52, 1162–1170

Brookes M. J., Hale J. R., Zumer J. M., Stevenson C. M., Francis S. T., Barnes G. R., et al. (2011). Measuring functional connectivity using MEG: methodology and comparison with fcMRI. Neuroimage 56, 1082–1104. 10.1016/j.neuroimage.2011.02.054

Buldú, J.M., Bajo, R., Maestú F, Castellanos N, Leyva I, Gil P, Sendiña-Nadal I, Almendral JA, Nevado A, del-Pozo F., 2011. Reorganization of functional networks in mild cognitive impairment. PLOS One. 6(5):e19584.

Buszaky, G. 2006. Rhythms of the Brain. Oxford University Press, New York 488 p.

Buzsaki, G. (2010). Neural syntax: Cell assemblies, synapsembles, and readers. Neuron, 68, 362–385.

Calhoun, V.D., Miller R, Pearlson G, Adali T., 2014. The chronnectome: time-varying connectivity networks as the next frontier in fMRI data discovery. Neuron 84(2), 262–274. doi:10.1016/j.neuron.2014.10.015

Calhoun, V.D. and T. Adalı., 2012. Multi-subject independent component analysis of fMRI: A decade of intrinsic networks, default mode, and neurodiagnostic discovery.IEEE Rev. Biomed. Eng.,5, 60–73.

Calhoun, V.D. and Adali, T. 2016. Time-Varying Brain Connectivity in fMRI Data: Whole-brain data-driven approaches for capturing and characterizing dynamic states. IEEE Signal processing Magazine 33:52–66 DOI:10.1109/MSP.2015.2478915

Candes, E.,, Romberg, J., Tao T., 2006. Stable signal recovery from incomplete and inaccurate measurements. Communications on Pure and Applied Mathematics 59, 1207–1223.

Canolty, R.T., Knight R.T., 2010. The functional role of cross-frequency coupling, Trends Cogn. Sci. 14,506–515.

Chang, C., Glover, G., 2010. Time–frequency dynamics of resting-state brain connectivity measured with fMRI. NeuroImage 50, 81–98

Colclough, G.L, Woolrich, M.W., Tewarie, P.K., Brookes, M.J., Quinn, A.J., Smith S.M., 2016. How reliable are MEG resting-state connectivity metrics? Neuroimage. 2016 Jun 1. pii: S1053–8119(16)30191-4. doi: 10.1016/j.neuroimage.2016.05.070. [Epub ahead of print]

Collins, F.S, Tabak, 2014. LA. Policy: NIH plans to enhance reproducibility. Nature. 30, 505 (7485):612–3.

Contreras, J. A., Goni, J., Risacher, S. L., Amico, E., Yoder, K., Dzemidzic, M., et al. (2017). Cognitive complaints in older adults at risk for Alzheimer’s disease are associated with altered resting-state networks. Alzheimer’s Dement. 6, 40–49. doi: 10.1016/j.dadm.2016.12.004

Cuesta, P.,, Garcés P., Castellanos NP, López ME, Aurtenetxe S, Bajo R, Pineda-Pardo JA, Bruña R, Marín AG, Delgado M, Barabash A, Ancín I, Cabranes JA, Fernandez A, Del Pozo F, Sancho M, Marcos A, Nakamura A, Maestú F.,2015. Influence of the APOE ε4 allele and mild cognitive impairment diagnosis in the disruption of the MEG resting state functional connectivity in sources space. J Alzheimers Dis. 44(2),493–505.

Cuesta, P., Barabash, A., Aurtenetxe, S., Garcés P, López ME, Bajo R, Llanero-Luque M, Ancín I, Cabranes JA, Marcos A, Sancho M, Nakamura A, Maestú F, Fernandez. A., 2014. Source Analysis of Spontaneous Magnetoencephalograpic Activity in Healthy Aging and Mild Cognitive Impairment: Influence of Apolipoprotein E Polymorphism. J. Alzheimers Dis. 43(1), 259–73

Damaraju, E., Allen, E., Belger A., Ford J., McEwen S., Mathalon D., Mueller B., Pearlson G., Potkin S., Preda A., et al., 2014. Dynamic functional connectivity analysis reveals transient states of dysconnectivity in schizophrenia. NeuroImage: Clinical 5,298–308.

Deco G, Corbetta M., 2011. The dynamical balance of the brain at rest. Neuroscientist. 17:107–123.

Delbeuck, X., Van der Linden M. Collette F., 2003. Alzheimer’s disease as a disconnection syndrome? Neuropsychol. Rev. 13(2),79–92.

Dimitriadis, S.I., Laskaris, N.A., Del Rio-Portilla Y, Koudounis G.C., 2009. Characterizing dynamic functional connectivity across sleep stages from EEG. Brain Topogr 22,119–133.

Dimitriadis, S.I., Laskaris, N.A., Tsirka, V., Vourkas, M., Micheloyannis S, Fotopoulos S (2010a): Tracking brain dynamics via time-dependent network analysis. Journal of Neuroscience Methods 193(1), 145–155.

Dimitriadis, S.I., Laskaris, N.A., Tsirka, V., Vourkas M, Micheloyannis S., 2010b. What does delta band tell us about cognitive Processes: a mental calculation study? Neuroscience Letters 483(1), 11–15.

Dimitriadis, S. I., Laskaris, N. A., Tsirka, V., Erimaki, S., Vourkas, M., Sifis, M., et al. (2011). A novel symbolization scheme for multichannel recordings with emphasis on phase information and its application to differentiate EEG activity from different mental tasks. Cogn. Neurodyn. 6, 107–113. doi: 10.1007/s11571-011-9186-5

Dimitriadis, S.I., Laskaris, N.A., Tsirka, V., Vourkas M, Micheloyannis S., 2012a. An EEG study of brain connectivity dynamics at the resting state. Nonlinear Dynamics, Psychology and Life Sciences 16(1), 5–22.

Dimitriadis, S.I., Kanatsouli, K., Laskaris, N.A., Tsirka V, Vourkas M, Micheloyannis, S., 2012b. Surface EEG shows that Functional Segregation via Phase Coupling contributes to the neural Substrate of Mental Calculations. Brain and Cognition 80(1), 45–52.

Dimitriadis, S.I., Laskaris, N.A., Tzelepi A, Economou G., 2012c. Analyzing Functional Brain Connectivity by means of Commute Times: a new approach and its application to track event-related dynamics. IEEE (TBE) Transactions on Biomedical Engineering 59(5), 1302–1309.

Dimitriadis, S.I., Laskaris, N.A., Simos, P.G., Micheloyannis S, Fletcher JM, Rezaie R, Papanicolaou, A.C., 2013a. Altered temporal correlations in resting-state connectivity fluctuations in children with reading difficulties detected via MEG. NeuroImage 83,307–331.

Dimitriadis, S.I., Laskaris, N.A., Tzelepi A., 2013b. On the quantization of time-varying phase synchrony patterns into distinct functional connectivity microstates (FCμstates) in a multi-trial visual ERP paradigm. Brain Topogr 26(3), 397–409.

Dimitriadis,.. SI, Sun, Yu, Kwok K, Laskaris NA, Thakor N, Bezerianos,A., 2013c. A tensorial approach to access cognitive workload related to mental arithmetic from EEG functional connectivity estimates. Conf Proc IEEE Eng Med Biol Soc 2940–2953.

Dimitriadis, S.I., Zouridakis, G., Rezaie R, Babajani-Feremi A, Papanicolaou A.C, 2015a. Functional connectivity changes detected with magnetoencephalography after mild traumatic brain injury. NeuroImage: Clinical 9,519–531.

Dimitriadis, SI, Laskaris NA, Micheloyannis S, 2015b. Transition Dynamics of EEG-based Network Microstates unmask developmental and task differences during mental arithmetic and resting wakefulness. Cogn Neurodyn 9(4), 371–387.

Dimitriadis, S.I., Sun, Yu., Kwok K, Laskaris NA, Thakor N, Bezerianos, A., 2015c. Cognitive Workload Assessment Based on the Tensorial Treatment of EEG Estimates of Cross-Frequency Phase Interactions. Ann Biomed Eng 43(4), 977–989.

Dimitriadis, S.I., Laskaris, N.A, Bitzidou MP, Tarnanas I, Tsolaki, M.N., 2015d. A novel biomarker of amnestic MCI based on dynamic cross-frequency coupling patterns during cognitive brain responses. Front Neurosci 9:350.

Dimitriadis, S.I., Sun, Y.,, Laskaris N, Thakor N, Bezerianos, A., 2016a. Revealing cross-frequency causal interactions during a mental arithmetic task through symbolic transfer entropy: a novel vector-quantization approach. IEEE Trans.Neural Syst.Rehabil Eng Oct;24(10),1017–1028.

Dimitriadis, S.I.,, Laskaris NA, Simos PG, Fletcher JM, Papanicolaou A.C, 2016b. Greater repertoire and temporal variability of cross-frequency coupling (CFC) modes in resting-state neuromagnetic recordings among children with reading difficulties. Front Hum Neurosci 10:163.

Dimitriadis, S. I., Salis, C., Tarnanas, I., and Linden, D. (2017a). Topological filtering of dynamic functional brain networks unfolds informative chronnectomics: a novel data-driven thresholding scheme based on orthogonal minimal spanning trees (OMSTs). Front. Neuroinform. 11:28. doi: 10.3389/fninf.2017.00028

Dimitriadis, S. I., and Salis, C. I. (2017b). Mining time-resolved functional brain graphs to an EEG-based chronnectomic brain aged index (CBAI). Front. Hum. Neurosci. 11:423. doi: 10.3389/fnhum.2017.00423

Dimitriadis, S. I., Antonakakis, M., Simos, P., Fletcher, J. M., and Papanicolaou, A. (2017c). Data-driven topological filtering based on orthogonal minimal spanning trees: application to multi-group MEG resting-state connectivity. Brain Connect. 7, 661–670. doi: 10.1089/brain.2017.0512

Dimitriadis, S. I., López, M. E., Bruña, R., Cuesta, P., Marcos, A., Maestú, F., et al. (2018a). How to build a functional connectomic biomarker for mild cognitive impairment from source reconstructed MEG resting-state activity: the combination of ROI representation and connectivity estimator matters. Front. Neurosci. 12:306. doi: 10.3389/fnins.2018.00306

Dimitriadis, S. I., Routley, B., Linden, D. E., & Singh, K. D. (2018b). Reliability of static and dynamic network metrics in the resting□state: A MEG□beamformed connectivity analysis.Front. Neurosci., 03 August 201| https://doi.org/10.3389/fnins.2018.00506

Dimitriadis SI (2018c). Complexity of brain activity and connectivity in functional neuroimaging.J Neurosci Res. 2018 Nov; 96(11):1741–1757. doi: 10.1002/jnr.24316.

Engel, A.K., Fries, P., Singer,W., 2001. Dynamic predictions: oscillations and synchrony in top-down processing. Nature reviews Neuroscience. 2,704–716.

Engel, A.K., Gerloff, C., Hilgetag CC, Nolte, G., 2013. Intrinsic coupling modes: Multiscale interactions in ongoing brain activity. Neuron 80,867–886.

Fischl, B., Salat, D.H., Busa E, Albert M, Dieterich M, Haselgrove C, van derKouwe A, Killiany R,Kennedy D, Klaveness S, Montillo A, Makris N, Rosen B, Dale, A.M., 2002. Whole Brain Segmentation. Neuron 33,341–355.

Florin, E., Baillet,. S, 2015. The brain’s resting-state activity is shaped by synchronized cross-frequency coupling of neural oscillations. Neuroimage. 111, 26–35.

Fox, M.D., Snyder, A.Z.,, Vincent JL, Corbetta M, Van Essen DC, Raichle, M.E., 2005. The human brain is intrinsically organized into dynamic, anticorrelated functional networks. Proc. Natl. Acad. Sci. U.S.A. 102, 9673–9678.

Fujisawa, S.,, Buzsaki, G., 2011. A 4 Hz oscillation adaptively synchronizes prefrontal, VTA, and hippocampal activities. Neuron. 72,153–165.

Gagniuc, P.A., 2017. Markov Chains: From Theory to Implementation and Experimentation. JohnWiley & Sons.

Gärtner, M., Brodbeck, V., Laufs, H., Schneider, G., 2015. A stochastic model for EEG microstates equence analysis. NeuroImage. 104, 199–208.

Gómez, C., Stam, C.J., Hornero R, Fernández A, Maestú, F., 2009. Disturbed beta band functional connectivity in patients with mild cognitive impairment: an MEG study. IEEE Trans Biomed Eng. 56(6),1683–90.

Gómez C., Hornero R., Abásolo D., Fernández A., Escudero J. (2009). Analysis of MEG background activity in Alzheimer’s disease using nonlinear methods and ANFIS. Ann. Biomed. Eng.37, 586–594. 10.1007/s10439-008-9633-6

Gómez C., Juan-Cruz C., Poza J., Ruiz-Gómez S. J., Gomez-Pilar J., Núñez P., et al. (2017). Alterations of effective connectivity patterns in mild cognitive impairment: an meg study. J. Alzheimers Dis. 10.3233/JAD-170475.

Gonzalez-Moreno A, Aurtenetxe S, Lopez-Garcia ME, del Pozo F, Maestu F, Nevado, A., 2014. Signal-to-noise ratio of the MEG signal after preprocessing. J. Neurosci. Methods. 222, 56–61.

Handwerker, D.A., Roopchansingh, V., Gonzalez-Castillo J, Bandettini, P.A., 2012. Periodic changes in fMRI connectivity. Neuroimage 63(3), 1712–9.

He, B.J., Snyder AZ, Zempel JM, Smyth MD, Raichle, M.E., 2008. Electrophysiological correlates of the brain’s intrinsic large-scale functional architecture. Proc. Natl. Acad. Sci. U.S.A. 105, 16039–16044.

Hipp, J.F., Hawellek DJ, Corbetta M, Siegel M, Engel, A.K, 2012. Large-scale cortical correlation structure of spontaneous oscillatory activity. Nat. Neurosci. 15:884–890.

Hutchison, R.M., Womelsdorf, T., Allen EA, Bandettini PA, Calhoun, V.D., Corbetta M, et al, 2013. Dynamic functional connectivity: promise, issues, and interpretations. Neuroimage 80,360–378.

Jack, C.R., Bennett,. DA., Blennow K, Carrillo MC, Feldman HH, Frisoni GB, Hampel H, Jagust WJ, Johnson KA, Knopman DS, Petersen RC, Scheltens P, Sperling RA, Dubois B, 2016. A/T/N: An unbiased descriptive classification scheme for Alzheimer disease biomarkers. Neurology. 2016 Jul 1. In Press

Jamal, W., Das, S., Oprescu, I.A., Maharatna, K., Apicella, F., Sicca, F., 2014. Classification of autism spectrum disorder using supervised learning of brain connectivity measures extracted from synchrostates. J. Neural. Eng. 11(4), 046019.

Jarvis, J.P., Shier, D.R., 1999. Graph-theoretic analysis of finite Markov chains. Appl. Math. Modeling Multidiscip. Approach, 85.

Jirsa, V., Muller, V., 2013. Cross-frequency coupling in real and virtual brain networks,” Front. Comput. Neurosci., 3,7–78.

Ioannides, A.A.,, Dimitriadis, S.I., Saridis GA, Voultsidou M, Poghosyan V, Liu L, Laskaris NA (2012):Source space analysis of event-related dynamic reorganization of brain networks Comput Math Methods Med (2012) 452503

Kirihara, K., Rissling, A.J. Swerdlow NR, Braff DL, Light, G.A, 2012. Hierarchical organization of gamma and theta oscillatory dynamics in schizophrenia. Biol Psychiatry. 15; 71(10),873–880.

Kitzbichler M.G., Smith, M.L., Christensen, S.R., Bullmore E., 2009. Broadband criticality of human brain network synchronization. PLoS Comput. Biol.S 5:e1000314

Kopell, N.J., Gritton,. H.J, Whittington MA, Kramer MA.Beyond the connectome: the dynome. Neuron. 17; 83(6), 1319–1328.

Laufs, H.,, Krakow,. K, Sterzer, P., Eger E, Beyerle A, Salek-Haddadi A, et al., 2003. Electroencephalographic signatures of attentional and cognitive default modes in spontaneous brain activity fluctuations at rest. Proc Natl Acad Sci U S A 100(19), 11053–11058.

Q.-H. Lin QH, J. Liu, Y.-R. Zheng, H. Liang, V. D.,(2013). Calhoun,.Semi-blind spatial ICA of fMRI using spatial constraints.Hum. Brain Mapp., 31(7), 1076–1088,

Lisman, J. E., & Idiart, M. A. (1995). Storage of 7 +/□ 2 short□term memories in oscillatory subcycles. Science, 267, 1512–1515.

Liu,X., Duyn JH., 2013. Time-varying functional network information extracted from brief instances of spontaneous brain activity. Proc Natl Acad Sci U S A 110(11).

López, M.E., Bruña, R., Aurtenetxe S, Pineda-Pardo JA, Marcos A, Arrazola J, Reinoso AI, Montejo P, Bajo R, Maestú, F., 2014. Alpha-band hypersynchronization in progressive mild cognitive impairment: a magnetoencephalography study.J. Neurosci., 34 (44), 14551–14559

López, M.E., Turrero, A., Cuesta P, López-Sanz D, Bruña R, Marcos A, Gil P, Yus M, Barabash A, Cabranes JA, Maestú F, Fernández, A., 2016. Searching for Primary Predictors of Conversion from Mild Cognitive Impairment to Alzheimer’s Disease: A Multivariate Follow-Up Study. J Alzheimers Dis. 5;52(1),133–143.

Maestú, F., Peña, J.M., Garcés P, González S, Bajo R, Anto Bagic A et al. 2015. A multicenter study of the early detection of synaptic dysfunction in Mild Cognitive Impairment using Magnetoencephalography-derived functional connectivity. Neuroimage Clin. 9, 103–109.

Mandal PK, Banerjee A, Tripathi M, Sharma A. A Comprehensive Review of Magnetoencephalography (MEG) Studies for Brain Functionality in Healthy Aging and Alzheimer’s Disease (AD). Front Comput Neurosci. 2018; 12: 60. doi: 10.3389/fncom.2018.00060

Mantini, D., Perrucci, M.G., Del Gratta C, Romani GL, Corbetta M., 2007.Electrophysiological signatures of resting state networks in the human brain. Proc Natl Acad Sci U S A 104(32), 13170–13175.

Martinetz, T. M., Berkovich, S. G., and Schulten, K. J. (1993). ‘Neural-gas’ network for vector quantization and its application to time-series prediction. IEEE Trans. Neural Netw. 4, 558–569. doi: 10.1109/72.238311

Hämäläinen MS, RJ Ilmoniemi. Interpreting magnetic fields of the brain: minimum norm estimates. Medical and Biological Engineering and Computing 32 (1), 35–42

Mylonas, D.S., Siettos, C.I., Evdokimidis I, Papanicolaou AC, Smyrnis,. N, 2015. Modular Patterns of Phase Desynchronization Networks During a Simple Visuomotor Task, Brain Topogr, 1–12.

Nacher, V., Ledberg, A., Deco, G., Romo, R., 2013. Coherent delta-band oscillations between cortical areas correlate with decision making. Proceedings of the National Academy of Sciences of the United States of America. 110, 15085–15090

Neymotin, S. A., Lazarewicz, M. T., Sherif, M., Contreras, D., Finkel, L. H., & Lytton, W. W. (2011). Ketamine disrupts θ modulation of γ in a computer model of hippocampus. Journal of Neuroscience, 31, 11733–11743.

Nolte, G. (2003).The magnetic lead field theorem in the quasi-static approximation and its use for magnetoencephalography forward calculation in realistic volume conductors. Phys. Med. Biol. 48, 3637–3652. doi: 10.1088/0031-9155/48/22/002

Nolte, G., Bai, O., Wheaton, L.,, Mari Z, Vorbach S, Hallett, M., 2004. Identifying true brain interaction from EEG data using the imaginary part of coherency. Clin Neurophysiol., 115, 2292–2307.

Nomi, J.S, Vij, S..G, Dajani DR, Steimke DR, Damaraju E, Rachakonda S, Calhoun, Uddin L.Q.,2016. Chronnectomic patterns and neural flexibility underlie executive function. Neuroimage http://dx.doi.org/10.1016/j.neuroimage.2016.10.026

Oostenveld, R., Fries, P., Maris, E., Schoffelen, J.M., 2011. FieldTrip: open source software for advanced analysis of MEG, EEG, and invasive electrophysiological data. Comput. Intell. Neurosci. 2011:156869.

Petersen, R.C., Negash, S., 2008. Mild cognitive impairment: an overview. CNS Spectr., 13,45–53.

Petersen, R.C., Roberts, R.O., Knopman, D.S., Boeve BF, Geda YE, Ivnik RJ, Smith GE, Jack Jr, C.R., 2009. Mild cognitive impairment: ten years later,” Arch.Neurol., 66, 1447–1455.

Petersen, RC., Doody, R., Kurz A, Mohs RC, Morris JC, Rabins PV, Ritchie K, Rossor M, Thal L,Winblad, B., 2001. Current concepts in mild cognitive impairment. Arch Neurol 58, 1985–1992.

Portet, F., Ousset, P.J., Visser, P.J., Frisoni GB, Nobili F, Scheltens P, Vellas B, Touchon J, and MCI Working Group of the European Consortium on Alzheimer’s Disease (EADC),2006. Mild cognitive impairment (MCI) in medical practice: A critical review of the concept and new diagnostic procedure. Report of the MCI Working Group of the European Consortium on Alzheimer’s Disease. J. Neurol. Neurosurg. Psychiatry 77, 714–718.

Poza J., Hornero R., Abásolo D., Fernández A., García M. (2007b). Extraction of spectral based measures from MEG background oscillations in Alzheimer’s disease. Med. Eng. Phys. Valladolid, 29, 1073–1083. 10.1016/j.medengphy.2006.11.006

Qiu, C., Kivipelto, M., von Strauss, E., 2009. Epidemiology of Alzheimer’s disease: occurrence, determinants, and strategies toward intervention. Dial. Clin. Neurosci. 11(2), 111–128.

Richiardi, J., Eryilmaz, H., Schwartz S, Vuilleumier P, Van De Ville D., 2011. Decoding brain states from fMRI connectivity graphs. NeuroImage, 56,616–626

Rosen, W.G.,Terry, R.D.,, Fuld PA, Katzman R, Peck, A., 1980. Pathological verification of ischemic score in differentiation ofdementias. Ann. Neurol. 7,486–488.

Sakoglu, U., G. D. Pearlson, K. A. Kiehl, Y. M. Wang, A. M. Michael, and V. D. Calhoun, 2010.A method for evaluating dynamic functional network connectivity and task-modulation: Application to schizophrenia,” Magn. Reson. Mater. Phys., 23, 351–366.

Sarvas, J. (1987). Basic mathematical and electromagnetic concepts of the biomagnetic inverse problem. Phys. Med. Biol. 32 11–22.

Schoffelen, J.M., Gross, J., 2009. Source Connectivity Analysis With MEG and EEG. Human Brain Mapping 30, 1857–1865.

Shirer, W.R., Ryali, S., Rykhlevskaia E, Menon V, Greicius, M.D., 2012. Decoding subject-driven cognitive states with whole-brain connectivity patterns. Cereb. Cortex 22,158–165.

Shuman DI, S. K. Narang, P. Frossard, A. Ortega, and P. Vandergheynst, “The emerging field of signal processing on graphs: Extending high-dimensional data analysis to networks and other irregular domains,” IEEE Signal Process. Mag., vol. 30, no. 3, pp. 83–98, May 2013.

Siegel, M., Donner, T.H., Engel, A.K., 2012. Spectral fingerprints of large-scale neuronal interactions. Nature reviews Neuroscience. 13,121–134.

Solin A, Jylänki P, Kauramäki J, Heskes T, van Gerven MAJ, Särkkä S. Regularizing solutions to the MEG inverse problem using space–time separable covariance functions. arXiv preprint. 2016; p. 1604.04931.

Stam, C.J.,, de Haan, W., Daffertshofer A, Jones BF, Manshanden I, van Cappellen van Walsum AM, Montez T, Verbunt JPa, de Munck JC, van Dijk, B.W., 2009. Graph theoretical analysis of magnetoencephalographic functional connectivity in Alzheimer’s disease. Brain. 132(1), 213–224.

Taulu, S., Simola, J., 2006. Spatiotemporal signal space separation method for rejecting nearby interference in MEG measurements. Phys Med Biol 51, 1759–1768.

Tewarie, P., Hillebrand, A., van Dijk, B. W., Stam, C. J., O’Neill, G. C., Van Mieghem, P., … Brookes, M. J. (2016). Integrating cross-frequency and within band functional networks in resting-state MEG: A multi-layer network approach. NeuroImage, 142, 324–336. http://doi.org/10.1016/j.neuroimage.2016.07.057

Toppi, J., Astolfi, L., Poudel, G. R, Innes CR, Babiloni F, Jones, R.D., 2015. Time-varying effective connectivity of the cortical neuroelectric activity associated with behavioural microsleeps. Neuroimage 124:421–432. doi: 10.1016/j.neuroimage.2015.08.059Yang CY, Lin, C.P., 2015. Time-Varying Network Measures in Resting and Task States Using Graph Theoretical Analysis. Brain Topogr. 28(4),529-408.

Van de Ville, D., Britz, J., Michel, C.M., 2010. EEG microstate sequences in healthy humans at rest reveal scale-free dynamics. Proc. Natl. Acad. Sci. U. S. A. 107(42), 18179–18184.

Van Veen, B. D., van Dronglen, W., Yuchtman, M., and Suzuki, A. (1997). Localization of brain electric activity via linearly constrained minimum variance spatial filtering. IEEE Trans. Biomed. Eng. 44, 867–880. doi: 10.1109/10.623056

van Wijk BC, Fitzgerald TH (2014) Thalamo-cortical cross-frequency coupling detected with MEG. Front Hum Neurosci 8:187

Wallis, G., Stokes, M., Cousijn, H., Woolrich, M., & Nobre, A. C. (2015). Frontoparietal and cingulo-opercular networks play dissociable roles in control of working memory. Journal of Cognitive Neuroscience, 27, 2019–2034.

Yu M., Engels M. M. A., Hillebrand A., van Straaten E. C. W., Gouw A. A., Teunissen C., et al. (2017). Selective impairment of hippocampus and posterior hub areas in Alzheimer’s disease: an MEG-based multiplex network study. Brain 140, 1466–1485. 10.1093/brain/awx050

