## Supplementary Material for "Modeling the Switching behavior of Functional Connectivity Microstates (FCμstates) as a Novel Biomarker for Mild Cognitive Impairment"

1. **The algorithmic steps for PAC estimation**
1. **Construction of the integrated Dynamic Functional Connectivity Graph**

There are numerous estimators to quantify the relationship between two oscillations with a different frequency profile. In the present study, we adopted the so-called phase-to-amplitude that quantifies the degree of modulation of **amplitude** of fast **frequency** by the **phase** of slow **frequency** employing the same formula of iPLV (PAC; Canolty and Knight, 2010; for further details see (Dimitriadis et al., 2015d, 2016b). S1 illustrates the algorithmic steps for the estimation of PAC between two time series oscillating in difference frequencies.

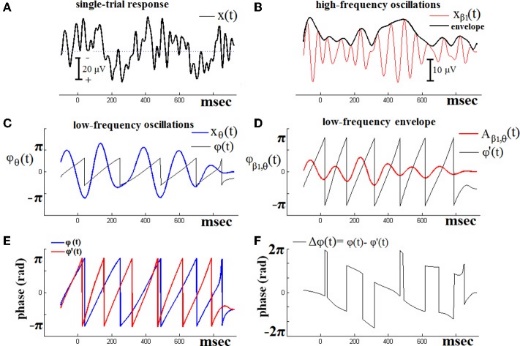

[**Figure 1**](https://www.ncbi.nlm.nih.gov/pmc/articles/PMC4611062/figure/F1/)

**The algorithmic steps for PAC estimation**. Using the first single-trial signal **(A)**, from the cognitive responses of a control subject, we demonstrate the detection of coupling between θ and β_1_ rhythm. To estimate θ-β_1_ PAC, the raw signal was band-pass filtered into both a **(B)** low-frequency θ (4–8 Hz) component where its envelope is extracted as well as **(C)** a high-frequency β_1_ (13–20 Hz) component where its instantaneous phase is extracted. **(D)** We then extracted the amplitude and the instantaneous phase of the band-passed β_1_ (13–20 Hz) and filtered this amplitude time series at the same frequency as θ (4–8 Hz), giving us the θ modulation in lower β amplitude. **(E)** We then extracted the instantaneous phase of both the θ-filtered signal and the θ-filtered lower-β amplitude and computed the phase-locking between these two signals. The latency depended differences **(F)**, will be used in estimating the phase-locking that will reflect the PAC-interaction between the two involved brain rhythms. This phase-locking represents the degree to which the lower β (β_1_) amplitude is comodulated with the θ phase.

The imaginary part of PLV is considered to be less susceptible to volume conduction e.ffects in assessing CFC interactions and was used in all subsequent analyses. While iPLV is not affected by volume conduction, it could be sensitive to changes in the angle between two signals, which do not necessarily imply a PLV change. In general, iPLV is only sensitive to non-zero-phase lags and is thus resistant to instantaneous self-interactions associated with volume conductance (Nolte et al., 2004).

2. **From Prominent Intrinsic Coupling Modes to Dominant Intrinsic Coupling Modes**

For each pair of 90 MEG sources, temporal segment and subject, the maximum PAC score was computed for each combination of *fφ* (frequency band of the phase modulators from δ to γ) and *fa* (frequency modulated oscillator ranging from 1 to 90 Hz in 1Hz steps). The maximum PAC value for each pair of sensors and temporal segment in both the amplitude and phase domain is given by the following equation:

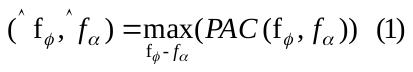

In this manner, we assessed whether the prominent CFC interaction (indexed by the maximum PAC value), and specifically the frequency of the low-frequency phase, was associated with the frequency of maximum power. If the two frequencies were identical this would imply that the observed prominent CFC interaction was driven by the power of the dominant frequency and not by its phase. In order to ensure that we did not include in further analysis CFC interactions of this type, we required a minimum frequency difference of 1Hz distance between the frequency of the low-frequency phase and that of the higher power, the one identified by equation 2 (maximum PAC value) and the one associated with the maximum spectral power, within the range of high amplitude.

After applying this criterion for the true PAC estimates based the significant iPLV strength of dominant type of interactions (either intra or PAC) for each pair of sources and across sliding windows were integrated within each of the eight frequency band, thereby yielding 28 possible pairwise PAC estimates among the eight frequency bands. iPLV has been adapted for the estimation of the eight intra-frequency interactions. For each participant the resulting ^TV^iPLV profiles constituted two 4D array of size [36 (intra and cross frequency pairs) x 2.401 (temporal segments) x 90 (sources) x 90 (sources)]. The first 4D array keeps the iPLV functional strength while the second 4D array stores the identity of prominent frequency pairs for every pair of sources at each temporal segment using one integer from 1 up to 36 for every possible coupling mode.

Complementary, we applied a previously defined surrogate analysis to reveal the dominant type of interaction for each pair of sources and at every snapshot of the ^TV^iPLV. We constructed 10,000 surrogate time‐series by cutting first at a single point at a random location within the middle of the original time series (between 1 min 40 sec and 2 min 20 sec over 4 mins of the whole duration of the resting-state) resulting to two temporal segments and then exchanging the order of the two resulting temporal segments (Dimitriadis & Salis, [2017](https://onlinelibrary.wiley.com/doi/full/10.1002/jnr.24316#jnr24316-bib-0029)b). Repeating this procedure leads to a set of surrogates with a minimal distortion of the original phase dynamics (see Dimitriadis & Salis, [2017](https://onlinelibrary.wiley.com/doi/full/10.1002/jnr.24316#jnr24316-bib-0029)b).

Finally, for each pair of sources and temporal segment, we estimated 1,000 iPLV surrogates for every pair of sources, temporal segments and possible intrinsic coupling modes (intra-frequency and cross-frequency interactions). Practically, we assigned a p‐value to each within and between frequencies interaction and for each pair of sources and temporal segments by comparing the original iPLV value with 1,000 surrogates iPLV^Surr^. Afterward, we corrected for multiple comparisons across 36 possible DICM in order to reveal a DICM per pair of sources and for each temporal segment independently for each subject. As a statistical threshold, we used (*p′* < *p*/36) where *p* = 0.05 in order to detect the DICM.

In total, there are three alternative scenarios:

a. no *p*‐value survived the multiple correction
b. more than one coupling mode survived and we finally selected the DICM with the maximum iPLV value or
c. only one survived the multiple correction

Finally, the FDR method (Benjamini and Hochberg, 1995) was employed to control for multiple comparisons by using p-values across all pair of sources within every snapshot of the DFCG. We set the expected proportion of false positives set to q ≤.01. The whole procedure of analysis is described in detail in (Dimitriadis et al., 2016b; Dimitriadis and Salis, 207b; Dimitriadis, 2018a,c). This procedure leads to two 3D arrays of dimensions [2.401 (temporal segments) x 90 (sources) x 90 (sources)] where the first keeps the iPLV coupling strength of the significant interactions and the second the dominant intrinsic coupling mode with an integer varying from 1 for δ up to 36 for γ_1_-γ_2_ CFC interaction.

**References**

Benjamini Y, Y HochbergControlling the false discovery rate—a practical and powerful approach to multiple testing.J R Stat Soc Ser B, 57 (1995), pp. 289-300

Dimitriadis, S.I.,, Laskaris NA, Simos PG, Fletcher JM, Papanicolaou A.C, 2016b. Greater repertoire and temporal variability of cross-frequency coupling (CFC) modes in resting-state neuromagnetic recordings among children with reading difficulties. Front Hum Neurosci 10:163.

Dimitriadis, S. I., and Salis, C. I. (2017b). Mining time-resolved functional brain graphs to an EEG-based chronnectomic brain aged index (CBAI). *Front. Hum. Neurosci.* 11:423. doi: 10.3389/fnhum.2017.00423

Dimitriadis, S. I., López, M. E., Bruña, R., Cuesta, P., Marcos, A., Maestú, F., et al. (2018a). How to build a functional connectomic biomarker for mild cognitive impairment from source reconstructed MEG resting-state activity: the combination of ROI representation and connectivity estimator matters. *Front. Neurosci.* 12:306. doi: 10.3389/fnins.2018.00306

Dimitriadis SI (2018c). Complexity of brain activity and connectivity in functional neuroimaging.[J Neurosci Res.](https://www.ncbi.nlm.nih.gov/pubmed/30259561) 2018 Nov; 96(11):1741-1757. doi: 10.1002/jnr.24316.
